# Exploring an EM-algorithm for banded regression in computational neuroscience

**DOI:** 10.1101/2023.09.22.558945

**Authors:** Søren A. Fuglsang, Kristoffer H. Madsen, Oula Puonti, Hartwig R. Siebner, Jens Hjortkjær

**Affiliations:** Danish Research Centre for Magnetic Resonance, Centre for Functional and Diagnostic Imaging and Research, Copenhagen University Hospital - Amager and Hvidovre, Copenhagen, Denmark; Hearing Systems Section, Department of Health Technology, Technical University of Denmark, Kgs. Lyngby, Denmark; Department of Applied Mathematics and Computer Science, Technical University of Denmark, Denmark, Kgs. Lyngby, Denmark; Martinos Center for Biomedical Imaging, Massachusetts General Hospital and Harvard Medical School, USA; Department of Neurology, Copenhagen University Hospital Bispebjerg, Copenhagen, Denmark; Institute for Clinical Medicine, Faculty of Health and Medical Sciences, University of Copenhagen, Copenhagen, Denmark

## Abstract

Regression is a principal tool for relating brain responses to stimuli or tasks in computational neuroscience. This often involves fitting linear models with predictors that can be divided into groups, such as distinct stimulus feature subsets in encoding models or features of different neural response channels in decoding models. When fitting such models, it can be relevant to allow differential shrinkage of the different groups of regression weights. Here, we explore a framework that allow for straightforward definition and estimation of such models. We present an expectation-maximization algorithm for tuning hyperparameters that control shrinkage of groups of weights. We highlight properties, limitations, and potential use-cases of the model using simulated data. Next, we explore the model in the context of a BOLD fMRI encoding analysis and an EEG decoding analysis. Finally, we discuss cases where the model can be useful and scenarios where regularization procedures complicate model interpretation.

## 1 Introduction

Understanding how the brain responds to different stimuli or tasks is a key objective in systems neuroscience. In recent years, significant progress has been made in developing models to analyze neural responses to complex, naturalistic stimuli (Hamilton and Huth, 2020). One potent approach is encoding models that combine stimulus features to predict channel-wise neural responses from fMRI or electrophysiology through linear regression (Dumoulin and Wandell, 2008; Kay et al., 2013; de Heer et al., 2017; Broderick et al., 2018; Di Liberto et al., 2015; Kell et al., 2018). Regularization is often needed in such models to reduce the risk of over-fitting and improve model prediction on held-out data. However, multiple - potentially co-linear - stimulus feature sets can complicate the assessment of the relative importance of the individual features. In this case, it can be relevant to allow for differential shrinkage of different groups of regression weights. For instance, neural responses to natural speech depend on both low-level acoustic features and higher-level phonetic or semantic features (Huth et al., 2016; Broderick et al., 2018; Di Liberto et al., 2015; Jain and Huth, 2018). Such feature groups vary on different time scales, have different dimensionality, and some features may not even be reflected in the neural response.

Several regularization strategies could have relevance for this situation (Massy, 1965; Hoerl and Kennard, 1970; Tikhonov and Arsenin, 1977; Golub et al., 1999; Sahani and Linden, 2002; Zou and Hastie, 2005; Tibshirani, 1996; Tibshirani et al., 2005; David et al., 2007; Park and Pillow, 2011). Recently, (Nunez-Elizalde et al., 2019; la Tour et al., 2022) proposed to divide predictors in encoding analyses into separate feature ”bands” (groups of predictors) and allow differential shrinkage of weights associated with these different bands. The authors proposed banded Ridge regression - also referred to as grouped Ridge regression (van de Wiel et al., 2021) - and showed that banded Ridge regression can yield higher out-of-sample predictive accuracy compared to standard Ridge regression in fMRI encoding analyses (Nunez-Elizalde et al., 2019; la Tour et al., 2022). Other methods that can incorporate grouping structure constraints and other priors (Yuan and Lin, 2006; Zhao et al., 2009; Friedman et al., 2010; Simon et al., 2013; Van De Wiel et al., 2016; Xu and Ghosh, 2015; Boss et al., 2023) could also be relevant in encoding analyses with grouped features.

Similar regression problems arise in decoding models, i.e., models that combine neural channels (voxels, electrodes) to predict a continuous stimulus- or task feature (Naselaris et al., 2011; Wu et al., 2006). One pertinent example is decoding models for EEG or MEG responses to continuous stimuli, like speech or music (Ding and Simon, 2012; O’Sullivan et al., 2014; Di Liberto et al., 2021). Here, regularized regression can be used to identify multidimensional finite impulse response (FIR) filters that predict stimulus features by combining different electrode signals at different time lags and/or different frequency bands (de Cheveigné et al., 2018; Mesgarani and Chang, 2012; Mirkovic et al., 2015). As with encoding models, it can be useful to allow for differential shrinkage of groups of weights corresponding to groups of predictors.

Here, we explore an empirical Bayes framework (Robbins, 1964; Efron and Morris, 1975, 1973; Morris, 1983) for estimating ”banded”-type regression models. The framework is closely related to automatic relevance determination (MacKay et al., 1994; Sahani and Linden, 2002; Park and Pillow, 2011; Wipf and Nagarajan, 2007; Tipping, 2001; Bishop, 2006) and can be seen as an extension of work by Tipping (2001). The framework can be used to formulate regularized regression estimators with different regularization properties. We use an expectation-maximization (EM) algorithm to tune hyperparameters that control variances of prior distributions. The model can be used to formulate priors that allow for differential shrinkage of groups of regression weights. The model can additionally be used to promote smoothness on groups of regression weights by encouraging correlation between related regression weights.

In this paper, we describe the modeling framework and provide simulations to illustrate the properties, limitations and potential use-cases of the described algorithm. Additionally, we illustrate properties of the model by using it to fit stimulus-response models on two publicly available datasets (Broderick et al., 2018; Nakai et al., 2022) and compare these model fits with the widely used Ridge estimator (Hoerl and Kennard, 1970). In the first dataset, we consider encoding models that use a set of audio features to predict BOLD fMRI responses to music (Nakai et al., 2022). In the second, we fit decoding models that use features of multi-channel time-lagged EEG data to predict an audio stimulus feature during a speech listening task (Broderick et al., 2018). Finally, we describe how simplifying model assumptions can allow for an efficient estimation in regression problems with multiple target variables. Software implementations of the model in both MATLAB and Python are available at https://github.com/safugl/embanded.

## 2 Regression model

Encoding and decoding analyses often involve linear regression models in the following form:

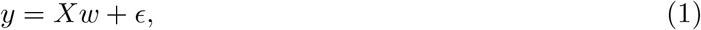

where *ϵ* is a zero-mean Gaussian variable.

Here, encoding models would treat stimulus- or task features as predictors, *X*, and a (preprocessed) neural response as target variable, *y*. Decoding models treat features extracted from neural activity measurements as predictors, *X*, and a stimulus- or task feature as target variable, *y*. The differences in direction of inference have large implications for model interpretation and model properties (Haufe et al., 2014; Naselaris et al., 2011) as well as how potential stimulus- or task confounders can be dealt with.

Despite their differences, such encoding and decoding models lead to regression problems that can be encompassed in a general formulation. Let *y* ∈ R^*M*×1^ denote a target variable where *M* indicates the number of observations. Moreover, let *X* denote a matrix with *D* predictors such that *X* ∈ R^*M*×*D*^. Suppose that *X* can be divided into a set of meaningful groups such that *X* = [*F*_1_, *F*_2_, …, *F*_*J*_], *j* = 1, 2, …, *j*, …*J*. Here, *F*_*j*_ denotes a group of *D*_*j*_ predictors, i.e., 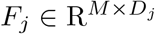. This group of predictors could, for example, be a stimulus feature subset in an encoding model, or response features of different voxels or electrodes in a decoding model. We thus consider a regression model in the following form (Nunez-Elizalde et al., 2019; la Tour et al., 2022):

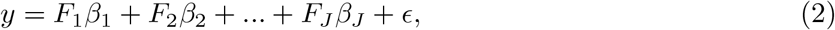

where *β*_*j*_ denotes regression weights for a given group of predictors, *F*_*j*_. Further, let *w* denote a column-vector collecting all the corresponding weights 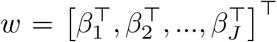, with *w* ∈ R^*D*×1^. We will assume that input variables and target variables have been centered and that no intercept term is needed in the regression model (since the intercept typically carries no meaningful information and hence shrinkage is typically not desired). The conditional distribution of the target variable *y* given a set of observed predictors *X* and given the model with parameters *w* and *ν* will be assumed to follow a Gaussian distribution in the following form:

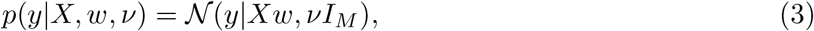

where 𝒩 (*y*|*Xw, νI*_*M*_) is used to indicate a multivariate Gaussian with mean *Xw* and a diagonal covariance matrix *νI*_*M*_ of size *M* × *M*. A zero-mean Gaussian prior distribution will be placed over the weights:

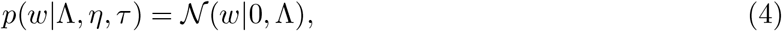

where Λ is a *D* × *D* block-diagonal covariance matrix. Such a prior distribution over the weights can be used to “shrink” the estimated regression weights. Here, we specifically define that Λ has a block structure defined as follows

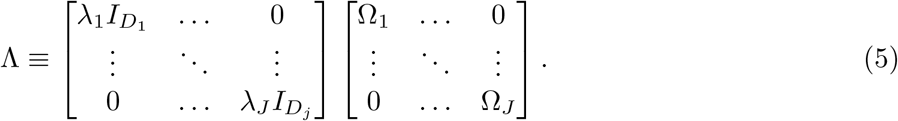

By introducing such a prior distribution over the weights, it is now possible to allow for differential shrinkage of different groups of weights. We define that the *j*-th block in Λ relates to a single group of predictors, *F*_*j*_, and that Ω_*j*_ has size *D*_*j*_ × *D*_*j*_. The column indices in *X* unique to predictor group *F*_*j*_ will thus index rows and columns in Λ for that block. The terms Ω_*j*_, *j* = 1, …, *J*, denote matrices that are assumed to be fixed a priori. For now, we will assume that they are identity matrices. The following parametrization (Rasmussen and Williams, 2005; Matérn, 1960) of these terms will later be used for encouraging local smoothness via a Matérn covariance function with order 3*/*2 and length scale *h*_*j*_:

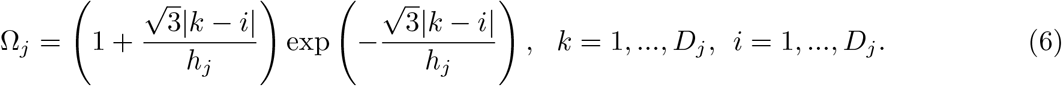

Note that rather than attempting to estimate *h*_*j*_ and impose hyperpriors on the *h*_*j*_ terms, we will fix these parameters a priori. We complete the model by placing Inverse-Gamma prior distributions over the *λ*_*j*_ terms and over the *ν* term:

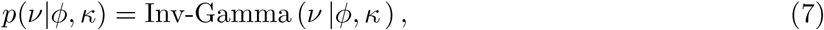

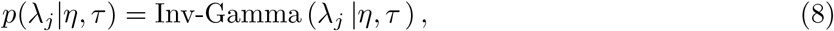

where Inv-Gamma (*x* | *a, b*) denotes an inverse-gamma distribution with shape *a*, scale *b* and mode *b*/(*a* + 1):

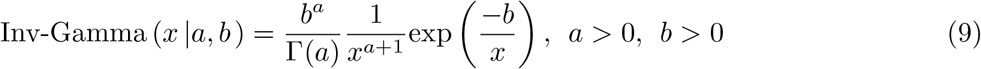

To simplify notation, we will henceforth let *λ* denote a set of 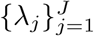 parameters and not highlight dependence on terms that remain fixed. We now need to regard *λ* and *ν* as random variables and by doing so the posterior distribution over all variables takes the form:

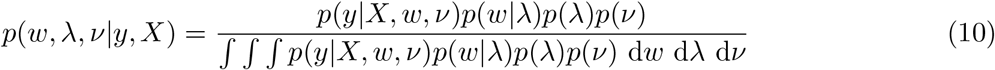

This expression cannot be evaluated analytically. In what follows, we will resort to an empirical Bayesian estimation procedure and assume that *p*(*w y, X*) =∫ ∫ *p*(*w, ν, λ X, y*) d*ν* d*λ* can be approximated by 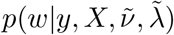 where 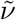 and 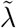 are fixed values obtained by maximizing the marginal posterior distribution:

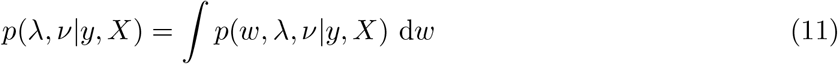

We will return to assumptions entailed by this procedure later. Our goal is to use an expectation-maximization (EM) framework (Dempster et al., 1977; Minka, 1998; Neal and Hinton, 1998) to maximize this marginal distribution. In brief, this involves lower-bounding ln *p*(*λ, ν y, X*) with a function ℱ(*λ, ν, q*) that is a functional of a valid probability distribution *q*(*w*) over *w*, and a function of the parameters *λ* and *ν* (Bishop, 2006):

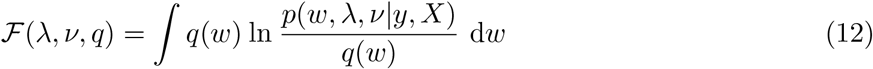

The framework involves alternating between an ”E-step” (finding a good lower bound) and an “M-step” (subsequently maximizing this bound) (Minka, 1998). Suppose we initialize the algorithm with *λ*^(*k*)^ and *ν*^(*k*)^ where *k* = 0. In the expectation step (E-step), we fix *λ* and *ν* to these values and maximize ℱ(*λ*^(*k*)^, *ν*^(*k*)^, *q*) with respect to *q*(*w*). It can be shown that the lower bound ℱ will touch the objective when *q*(*w*) = *p*(*w y, X, λ*^(*k*)^, *ν*^(*k*)^) *q*^(*k*)^ where the posterior distribution of *w* given a fixed set of parameters *λ* and *ν* takes the form:

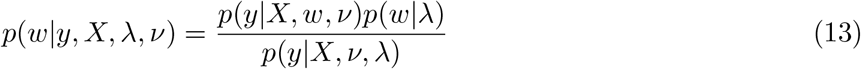

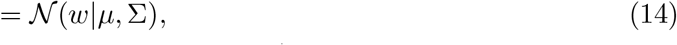

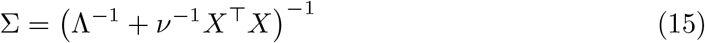

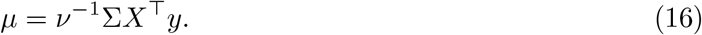

We will henceforth let 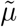 and 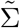 denote mean and covariance of *p*(*w*|*y, X, λ*^(*k*)^, *ν*^(*k*)^). Defining *q*(*w*) given *λ*^(*k*)^ and *ν*^(*k*)^ is the E-step. In the maximization step (M-step) we keep this distribution fixed and now maximize ℱ (*λ, ν, q*^(*k*)^) with respect to *λ* and *ν*. Ignoring irrelevant terms, one can show that we need to find a set of parameters *λ*^(*k*+1)^ and *ν*^(*k*+1)^ that maximizes the following expression:

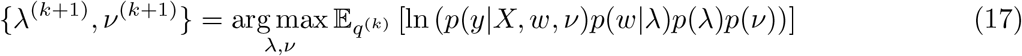

where 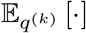 denotes expectation under *q*^(*k*)^ and where

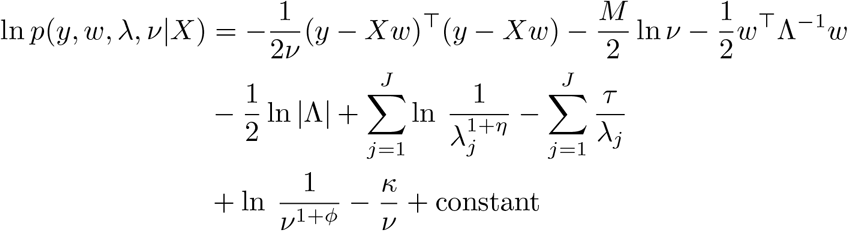

Recall that *w* is a column vector that contains all *β*_*j*_ weights and that Λ is a block diagonal matrix. We know that *p*(*w*|*y, X, λ*^(*k*)^, *ν*^(*k*)^) is Gaussian with mean 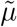 and covariance 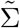. Let 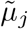 denote blocks in 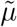 associated with predictor group *j*. Further, let 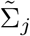 denote a block of size *D*_*j*_ × *D*_*j*_ along the diagonal in 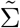 associated with predictor group *j*. We again discard irrelevant terms, and now see that the M-step amounts to maximizing the following expression with respect to *ν* and *λ*:

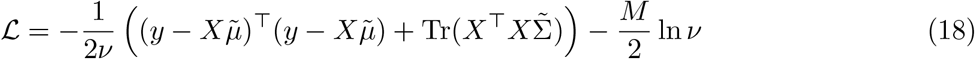

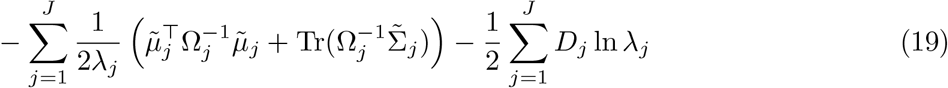

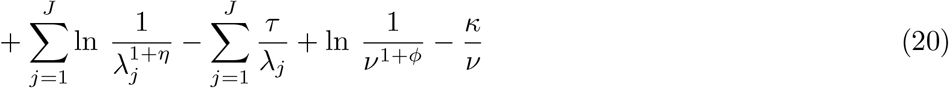

Finding a set of *λ* and *ν* parameters that fullfil *∂ ℒ/∂λ*_*j*_ = 0 and *∂ ℒ/∂ν* = 0 yields the following closed-form update rules:

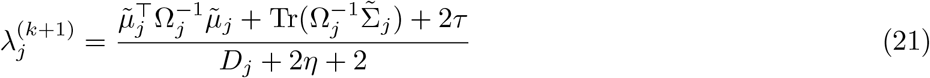

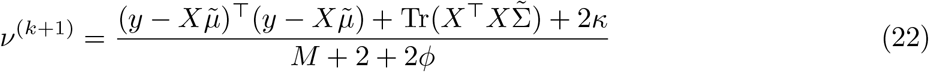

The procedure of fitting the model thus involves (I) specifying hyperprior parameters *ϕ, κ, η* and *τ*, (II) starting with an initial guess for *λ*_1_, *λ*_2_, …*λ*_*J*_ and *ν*, (III) estimating 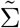 and 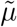, (IV) updating *λ*_*j*_ and *ν* according to Eq. 21 and 22, and subsequently iterating between step (III) and (IV) until convergence.

### 2.1 Predictions

The above procedure yields a set of hyperparameters 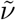 and 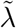 from the training data that may be sensible in several circumstances, and we will consequently focus on 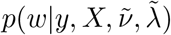. For predictive purposes, we assume that the following approximation holds:

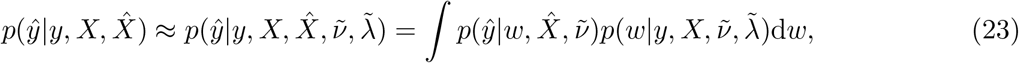

That is, to make predictions about a novel response variable, *ŷ*, given a new set of predictors 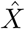 and inferred model parameters, we focus on 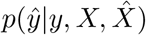 and let *p*(*λ, ν | X, y*) be represented by a delta function 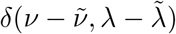 (Tipping, 2001). The resulting distribution is Gaussian with mean 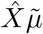 and covariance 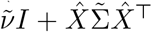. The terms 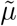 and 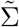 here refer to the estimates in Eq. 15 and Eq. 16 respectively. To make point predictions, we may now focus on the estimated model weights 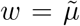 and simply compute 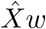. Our focus will be on such point predictions.

For the interested reader and illustrative purposes, we visualize variability in 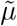 in Figure 1-3 based on Eq. 14. It is important to stress that 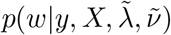 is conditional on the point estimates of 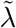 and 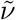 and that it may fail to be even close to a reasonable approximation of ∫ ∫ *p*(*w, ν, λ*|*X, y*) d*ν* d*λ* (Wolpert and Strauss, 1996).

**Figure 1.**
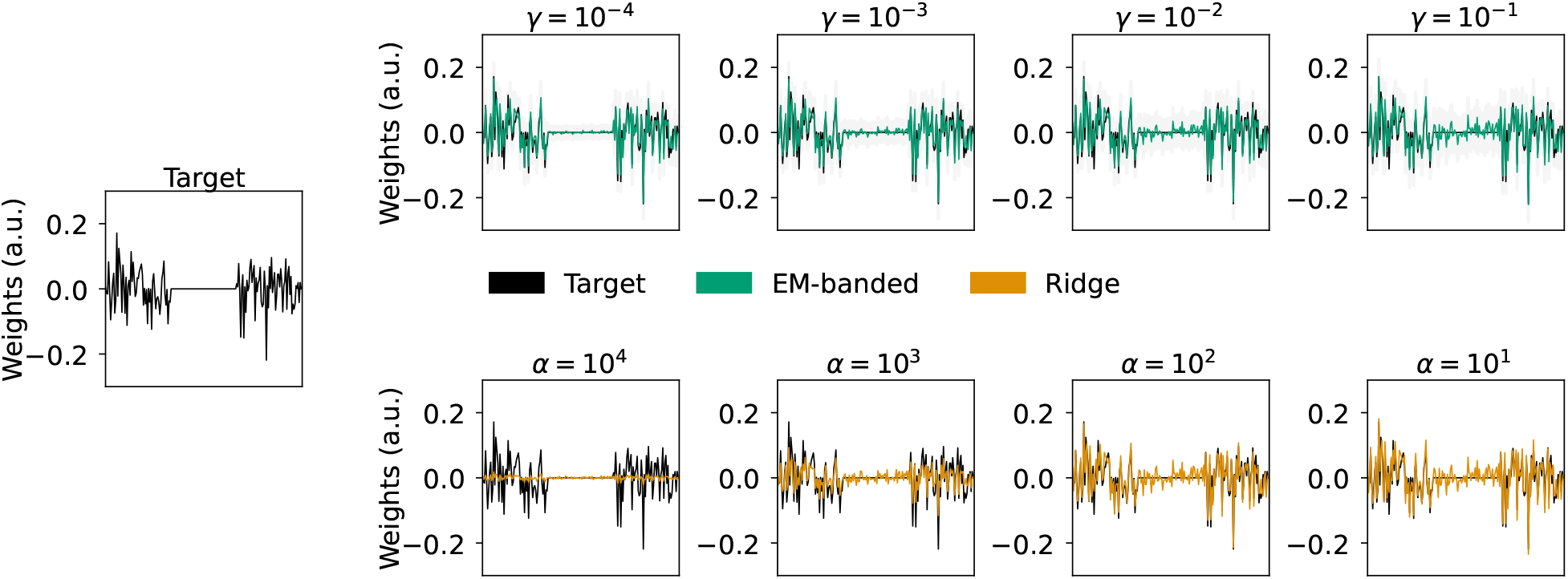
Simulation 1 illustrating regularization properties in a regression problem with three groups of predictors. Left middle panel: simulated target weights (black). The predictors are divided into three distinct predictor groups, *F*_1_, *F*_2_ and *F*_3_. Target weights associated with *F*_2_ are equal to zero. Top row: weights estimated with EM-banded model for different values of *η* = *ϕ* = *τ* = *κ* = *γ*. Shaded gray errorbars depict 1% respectively 99% percentile of samples from a multivariate Gaussian with mean and covariance defined according to Eq. 14. Bottom: weights estimated with Ridge regression model for different Ridge regularization parameters *α*. The target weights are shown in black.

**Figure 2.**
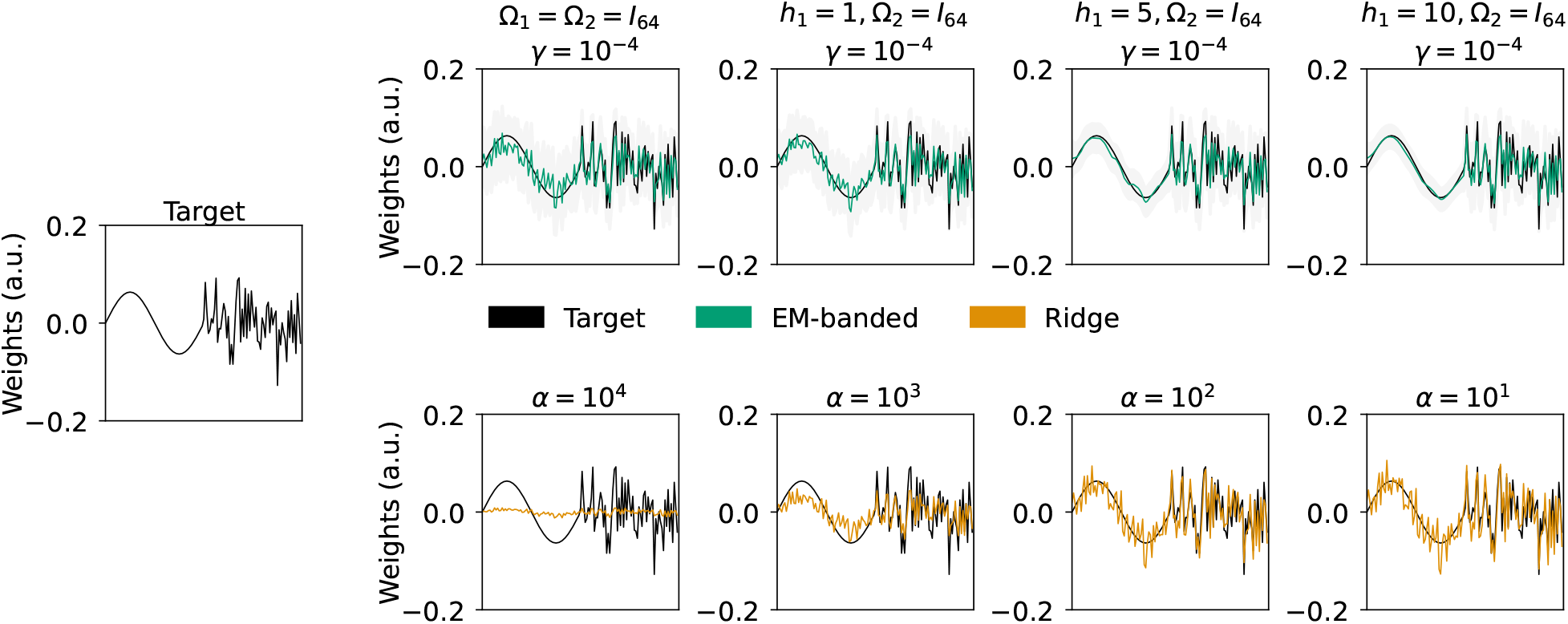
Simulation 2 illustrating effects of smoothness encouragement. Left middle panel: simulated target weights (black) divided into two distinct predictor groups, *F*_1_ and *F*_2_. Weights associated with *F*_1_ are sinusoidal, while weights associated with *F*_2_ are drawn from a Gaussian. Top row: weights estimated with the EM-banded model for *γ* = 10^−4^ but with different parameterizations for Ω_1_ (defined in Eq. 6). Specifically, we change the *h*_1_ term in this equation in order to encourage smoothness. Shaded gray errorbars depict 1% respectively 99% percentile of samples from a multivariate Gaussian with mean and covariance defined according to Eq. 14. Bottom: weights estimated with Ridge regression model for different Ridge regularization parameters *α*. The target weights are shown in black.

**Figure 3.**
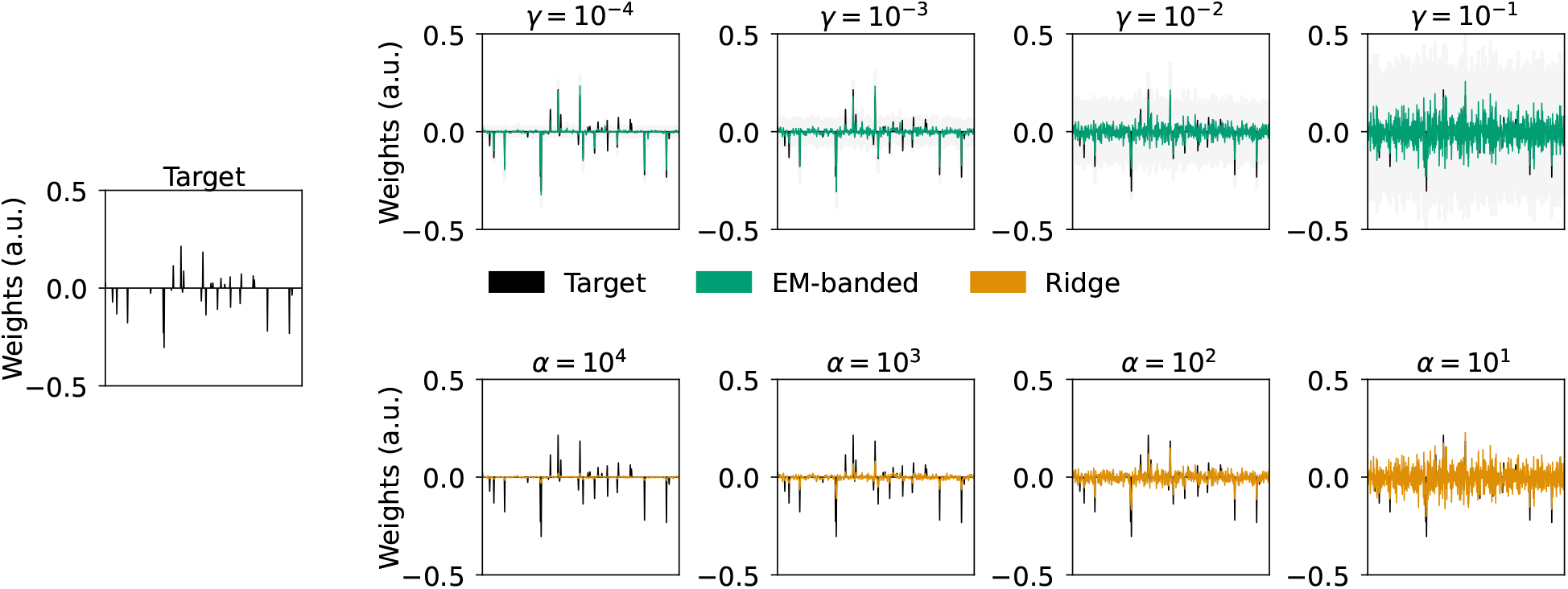
Simulation 3 illustrating differential shrinkage of individual weights. Left middle panel: simulated target weights (black). Top row: weights estimated with EM-banded model for different values of *η* = *ϕ* = *τ* = *κ* = *γ*. Shaded gray errorbars depict 1% respectively 99% percentile of samples from a multivariate Gaussian with mean and covariance defined according to Eq. 14. Bottom: weights estimated with Ridge regression model for different Ridge regularization parameters *α*. The target weights are shown in black.

## 3 Simulations

In addition to evaluating the predictive accuracy of encoding- or decoding models on held-out data, neuroimaging studies often inspect and interpret point estimates of model parameters. In linearized encoding models, model weights are often assumed to be directly interpretable in terms of the underlying brain activity generating a measured response (Haufe et al., 2014). Weights of general linear models (GLMs) in fMRI analysis (Friston et al., 1994) or temporal response functions in M/EEG analysis (Ding and Simon, 2012; Lalor et al., 2009) are prominent examples. However, the choice of regularization can significantly impact the relative magnitude of the estimated weights, in particular in problems with many, potentially correlated, sets of predictors. It can therefore be relevant to simulate data with known target model weights in order to better understand the properties of regularized regression estimators and identify situations where they can be useful or can complicate interpretation.

Below, we consider different simulations to illustrate the properties of the described model. We compare the proposed EM-based banded regression estimator (henceforth EM-banded) with a standard Ridge regression estimator: *w* = (*X*^*T*^ *X* + *αI*)^−1^ *X*^*T*^ *y* (Hoerl and Kennard, 1970). To simplify readability we consider simulations where we fix the parameters related to the Inverse-Gamma priors to the same value, *γ*, such that *τ* = *η* = *κ* = *ϕ* = *γ*. Notice that the hyperpriors become broad when *γ* approaches zero.

### 3.1 Simulation 1: suppressing contributions from a group of predictors

In models with groups of predictors representing different stimulus feature spaces, it is sometimes needed to identify entire feature spaces that can be excluded from an analysis without compromising predictive accuracy. To simulate such a situation, we here define three stimulus feature spaces, *F*_1_, *F*_2_ and *F*_3_, and one response variable *y* which contains mixed versions of *F*_1_ and *F*_3_, but not *F*_2_ (*F*_2_ does not contribute to the response). We thus simulate the response as *y* = *F*_1_*w*_1_ + *F*_3_*w*_3_ + *ϵ* where *ϵ* is a Gaussian noise term. In this situation, we thus need the estimator to identify (close to) zero weights for *F*_2_. Each feature space has 64 predictors and a total of *M* = 1024 observations are simulated. The weights, *w*_1_ and *w*_3_, as well as the feature space *F*_1_ and *F*_3_ are drawn from Gaussian distributions. The mixed target signal and the noise term have equal power. We compare weights estimated by the EM-banded model with weights estimated by Ridge regression models. To exemplify how hyperparameters controlling inverse-gamma hyperpriors can impact the estimated weights, we fit models for *γ* fixed to 10^−4^, 10^−3^, 10^−2^ and 10^−1^. The Ridge *α* hyperparameter is set to 10^4^, 10^3^, 10^2^ and 10^1^. The terms Ω_1_, Ω_2_ and Ω_3_ are identity matrices.

Figure 1 shows results from Simulation 1. The Ridge regression model yields ”dense” solutions for all considered Ridge hyperparameters, *α*, with no dramatic overall scale differences between weights associated with *F*_1_, *F*_2_ or *F*_3_. In contrast, the EM-banded model excessively shrinks weights associated with feature set *F*_2_ for smaller values of *γ*. As expected, the EM-banded model tends to show signs of under-shrinkage when the inverse-gamma prior distributions have little mass near zero and in these cases, the overall scale differences between the different sets of weights associated with *F*_1_, *F*_2_ and *F*_3_ are smaller.

### 3.2 Simulation 2: encouraging smoothness for a feature subset

It is sometimes relevant to encourage smoothness on sets of weights, for instance, in analyses that approximate smooth FIR filters. An example hereof is fMRI encoding analyses that model the relation between a task-regressor augmented with multiple time-lags and the BOLD voxel response. In this case, it can *a priori* be expected that the approximated FIR filter should be smooth due to hemodynamic properties of the BOLD signal (Goutte et al., 2000). It can similarly be relevant to assume smoothness across spectral channels in analyses that model relations between average stimulus power spectra and BOLD fMRI responses.

Here, we explore one way of encouraging smoothness using the definition of Ω_*j*_ in Eq. 6. To illustrate the effect of Ω_*j*_, we consider a simulated regression problem with two feature spaces, *F*_1_ and *F*_2_. Each feature space has 64 predictors and is drawn from a Gaussian distribution. Data are simulated as *y* = *F*_1_*w*_1_ +*F*_2_*w*_2_ +*ϵ* where *ϵ* is drawn from a Gaussian. A total of *M* = 1024 observations are simulated. The weights associated with the first feature space *w*_1_ are simulated to be sinusoidal (i.e., smooth), while *w*_2_ weights are Gaussian. The signal-to-noise ratio (SNR) is -5 decibels (dB) and we fit the EM-banded models with *γ* fixed to a low value of 10^−4^ (i.e., corresponding to a broad hyperprior). The Ω_1_ term associated with *F*_1_ is parameterized according to Eq. 6 with *h*_1_ set to 1, 5, and 10 to illustrate its effect.

Figure 2 shows weights estimated with Ridge estimators for different *α* values (below) and weights estimated by the EM-banded estimators (above). At higher levels of *h*_1_, the EM-banded approximates the weight distribution of the first feature space *F*_1_ with high accuracy. The Ridge estimator, on the other hand, yields weights that tend to show more rapid (high frequency) fluctuations across neighboring weights associated with *F*_1_ compared to the target weights.

### 3.3 Simulation 3: encouraging differential shrinkage of individual weights

In simulation 1, the EM-banded model was used to shrink weights of an entire feature space (i.e., groups of predictors). The model can similarly encourage differential shrinkage of individual weights. To illustrate this, we consider a simulation where we assume that the number of groups equals the number of predictors (i.e., each group contains a single predictor). We simulate 512 predictors, *F*_1_, *F*_2_, …, *F*_512_, and a response variable *y* which contains mixed versions of some of these predictors. A total of *M* = 1024 observations are simulated. It is assumed that *y* = ∑_*j*_ *F*_*j*_*w*_*j*_ + *ϵ* where *ϵ* is Gaussian noise. We assume that a high proportion of the target weights *w*_*j*_ are equal to zero. Each row in [*F*_1_, *F*_2_, …, *F*_512_] is drawn from a multivariate Gaussian with zero mean and a covariance defined as *C* = exp(−0.3 |*k* – *i*| ^2^), *k* = 1, …, 512, *i* = 1, …, 512. The SNR is equal to 0 dB. We again compare Ridge regression model fits with EM-banded model fits using the same set of fixed hyperparameters as in simulation 1.

The estimated weights for the Ridge and EM-banded model are shown in figure 3. It is evident that the EM-banded estimator with *γ* = 10^−4^ excessively shrinks several of the individual weights. In effect, the EM-banded model recovers the target weights with relatively high accuracy. In comparison, the Ridge estimator tends to dilute effects across neighboring (correlated) predictors and pull weights towards zero when *α* attains high values.

### 3.4 Simulation 4: Interpretation of weights with correlated predictors

Until now, we have not explored how strong correlations among groups of predictors can impact results. Models involving correlated groups of predictors occur often in EEG or fMRI encoding models that model responses to naturalistic stimuli (Hamilton and Huth, 2020). To exemplify this situation, this simulation considers the example of predicting responses to natural speech. Speech signals can be characterized by their lower-level acoustic features as well as higher-level phonetic or semantic features (Huth et al., 2016; Broderick et al., 2019). The different hierarchical levels can each be modeled by multidimensional features, but the statistics of natural speech imply a degree of correlation between such feature groups (see e.g., panel B in figure 4). This can complicate inferences about neural responses unique to a given group.

**Figure 4.**
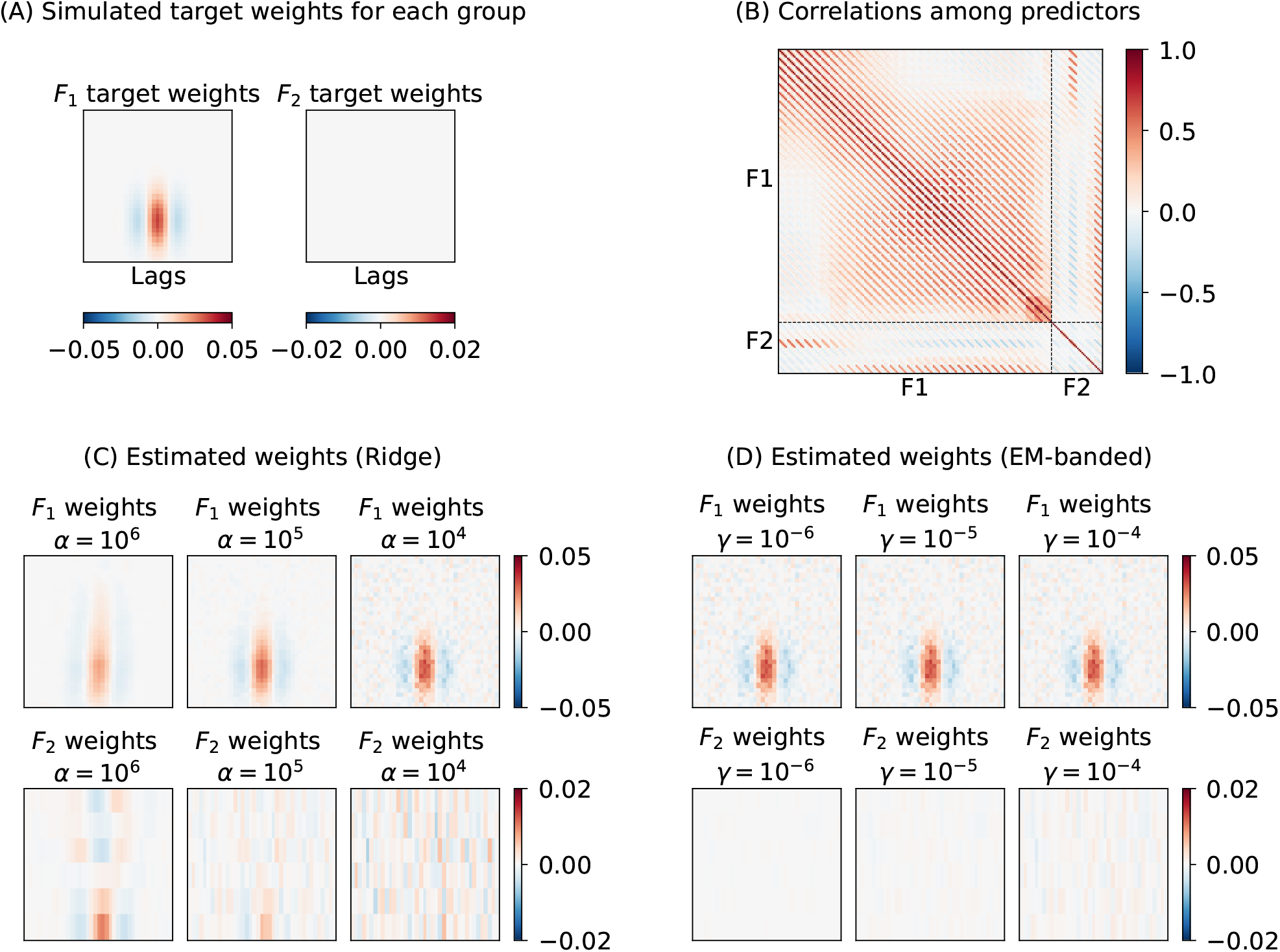
Simulation 4 illustrating behavior of the Ridge estimator and the EM-banded estimators when there are strong correlations among groups of predictors, *F*_1_ and *F*_2_. The target variable is simulated as *y* = *F*_1_*w*_1_ + *ϵ*. (A) Target weights associated with each of the two feature sets. (B) Matrix of Pearson correlation coefficients between predictors. Dashed lines are used to highlight predictor groups, *F*_1_ and *F*_2_. (C) Weights estimated with Ridge estimator for each of the two groups. (D) Weights estimated with the EM-banded estimator for each of the two groups. Ridge estimators with *α* set to 10^6^, 10^5^ and 10^4^ are shown. EM-banded estimators were considered with *γ* set to 10^−6^, 10^−5^ and 10^−4^

To illustrate this, we simulate responses to speech acoustic and phonetic features extracted from natural speech audio (Garofolo et al., 1993) (see section 4 in the Supplementary Material for details on feature extraction). A first feature set *F*_1_ contains time-lagged spectrogram features and *F*_2_ contains time-lagged regressors created from phonetic annotations. We simulate a response as *y* = *F*_1_*w*_1_ + *ϵ*, i.e., only the feature set associated with the spectrogram features *F*_1_ affects the response variable, *y*. The SNR is equal to 0 dB and the target filter weights, *w*_1_, are shown in figure 4A. We again compare Ridge and EM-banded regression model fits using a set of fixed hyperparameters.

Figure 4 shows the results of this simulation. The EM-banded estimator accurately recovers the weights associated with the speech spectrogram *F*_1_ while excessively shrinking the weights associated with the phonetic features *F*_2_, as desired. The Ridge estimator, on the other hand, pulls weights associated with correlated predictors in the two feature groups towards each other at higher levels of regularization. This has the undesired consequence that a temporally located response function associated with the phonetic feature group *F*_2_ emerges. This can complicate the interpretation of the weights associated with *F*_2_, e.g., if such point estimates of weights were interpreted or further analysed in second-level inferences. Simulating the opposite case, i.e., simulating only responses to phonetic features (and no contribution from spectrogram features), leads to the equivalent result: weights associated with the spectrogram feature group show temporally located response functions (see section 4 in the Supplementary Material). The described effect depends on the degree of correlation among predictors and on the amount of data available for fitting the models. We note that high levels of regularization often occur in practice for Ridge estimators, for instance, when tuning the regularization term to maximize the Pearson correlation coefficient between model predictions and target variables (due to its insensitivity to scaling and shift mismatches). For a related, but more stylized simulation, see section 5 in the Supplementary Material.

This simulation highlights a potential source of pitfall for the interpretation of model weights in situations with co-varying features affecting the response. To illustrate this further, consider again the simulation where the response is driven by the first feature set *F*_1_, but now only the correlated set *F*_2_ is included in the model. Figure 5 depicts the estimated weights by Ridge models in this case. The model with only *F*_2_ predictors now misleadingly indicates a temporally located response for all of the considered *α* parameters, despite the fact that the target regression weights associated with *F*_2_ are zero. Interpretation of the reduced model is clearly different from the model that includes both groups of predictors.

**Figure 5.**
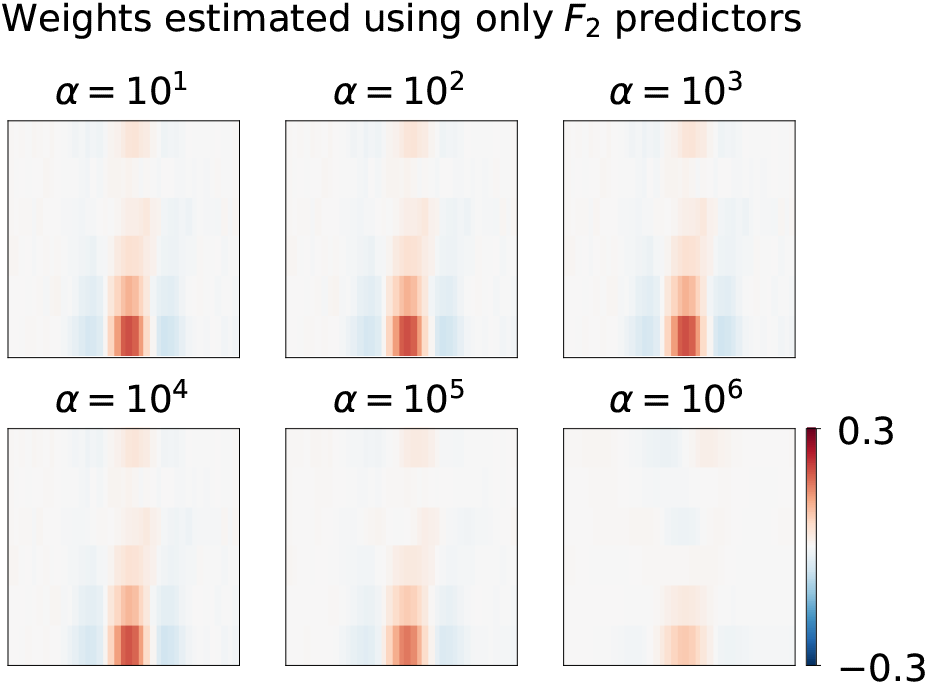
Overlooking important features in a Ridge regularized encoding model. The outcome variable is again simulated as *y* = *F*_1_*w*_1_ + *ϵ*, i.e., the simulated target is simulated to be affected by *F*_1_ and not *F*_2_, but only *F*_2_ is included in the prediction model. Panels reflect estimates with different Ridge regularization strengths.

### 3.5 Simulation 5: visualizing behavior under unfavorable signal-to-noise ratios

As illustrated in Simulation 3, the EM-banded estimator can be used to encourage differential shrinkage of weights associated with individual predictors. The behavior of such an estimator will depend on several factors, including the amount of data available to support the identification of *λ*, correlations among predictors, as well as the SNR. Here, we illustrate the behavior of the EM-banded estimator in a simulation with poor SNR. We simulate two predictors *F*_1_ and *F*_2_ and a response variable *y* = *F*_1_*w*_1_ + *F*_2_*w*_2_ + *ϵ*. The predictors as well as the noise term *ϵ* are drawn from Gaussian distributions. The target weights are fixed to non-zero values. The SNR is approximately − 20 dB. We consider simulations where the number of observations is 512, 1024 and 8192, and we fit EM-banded models to the data from each simulation. We focus on estimators with *γ* = 10^−4^ respectively *γ* = 10^−2^ to illustrate how differences in these prior choices affect the estimated parameters. This procedure is repeated many times to elucidate the effect.

Figure 6 shows 2D histogram visualizations of the estimated weights *w*_1_ and *w*_2_ from these simulations. Notably, the estimator with the lower *γ* = 10^−4^ tends to exhibit excessive shrinkage to either of the two weights when the number of observations are low (*M* = 512). Such over-shrinkage is undesirable. The estimator even collapses to *w*_1_ ≈*w*_2_ ≈0 occasionally, which again is undesirable. Increasing *γ* to 10^−2^ here yields dramatically different estimates and less differential shrinkage of the two weights (increasing *γ* encourages solutions that allow for less excessive shrinkage). These simulations underline subtleties that can yield dramatic effects on the estimated weights, which is important to keep in mind when focusing on ”default” priors for *λ*_*j*_.

**Figure 6.**
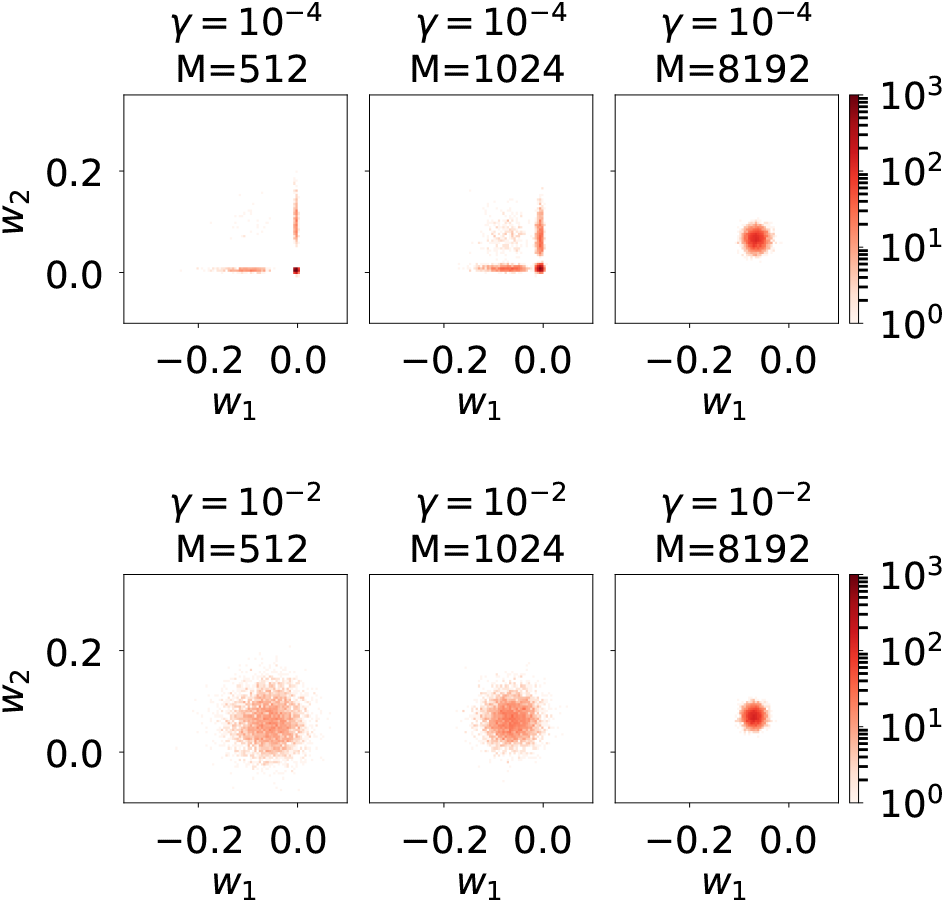
Simulation 5 illustrating regularization properties with only two predictors and a poor SNR. 2D histogram visualizations of estimated regression weights, *w*_1_ and *w*_2_. Columns show simulations with increasing number of observations. Top row: weights estimated with the EM-banded estimator with *γ* = 10^−4^. Bottom row: weights estimated with the EM-banded estimator with *γ* = 10^−2^.

## 4 fMRI encoding analysis with ”stimulus-irrelevant” predictors

Building encoding models often involve deciding on which of multiple potentially relevant feature sets to include as predictors. Choosing features both implies a risk of overlooking feature sets that are potentially relevant, as well as including feature sets that are irrelevant for predicting the target variables. In this section, we investigate how the inclusion of stimulus-irrelevant predictors - here defined as simulated noise predictors that are unrelated to the task or stimuli - can impact results in an fMRI encoding analysis and how the EM-banded estimator may help suppress their contribution.

The example uses a publicly available BOLD fMRI dataset that has been described in Nakai et al. (2022, 2021). The dataset contains BOLD fMRI data acquired from five participants listening to excerpts from music in 12 training runs and 6 test runs. Each functional run lasted 10 min and consisted of 40 musical clips that each were 15 s long. We focused on encoding models with three feature sets, *F*_1_, *F*_2_ and *F*_3_. The first two feature sets, *F*_1_ and *F*_2_, were related to the music stimuli: a 10-dimensional genre-label feature given by Tzanetakis and Cook (2002) indicating which of 10 musical genres a given stimulus is assigned to (*F*_1_), and a time-averaged 32-dimensional spectrogram of the audio stimulus (*F*_2_). As a the third feature set (*F*_3_) unrelated to the stimuli, we included a 400-dimensional multivariate Gaussian noise. *F*_1_, *F*_2_ and *F*_3_ were considered separate groups for the EM-banded model. For simplicity, we always initialize the EM algorithm with *λ*_*j*_ and *ν* parameters set to 1 and define that no more than 200 iterations should be considered. It may sometimes be advisable to use different initialization strategies. More details related to fMRI preprocessing, audio feature extraction, and model fitting are described in section 1 in the Supplementary Material.

Figure 7 shows the predictive accuracies obtained with the Ridge estimator and the EM-banded estimator. Predictive accuracy is here evaluated both as Pearson correlation coefficient between the predicted and measured BOLD fMRI response data, and as explained variance ratio. Compared to the Ridge estimator, the EM-banded estimator yielded higher predictive accuracy for all subjects in both metrics (all *p <* 0.05, Wilcoxon signed-rank tests across all voxels used for fitting models). The improvement in terms of Pearson correlation coefficient among the top 10000 voxels was (mean ± standard deviation): 0.0335 ± 0.0515 (sub-001), 0.0411 ±0.0629 (sub-002), 0.0568 ±0.0699 (sub-003), 0.0391 ±0.0637 (sub-004), 0.0298 ± 0.0522 (sub-005).

**Figure 7.**
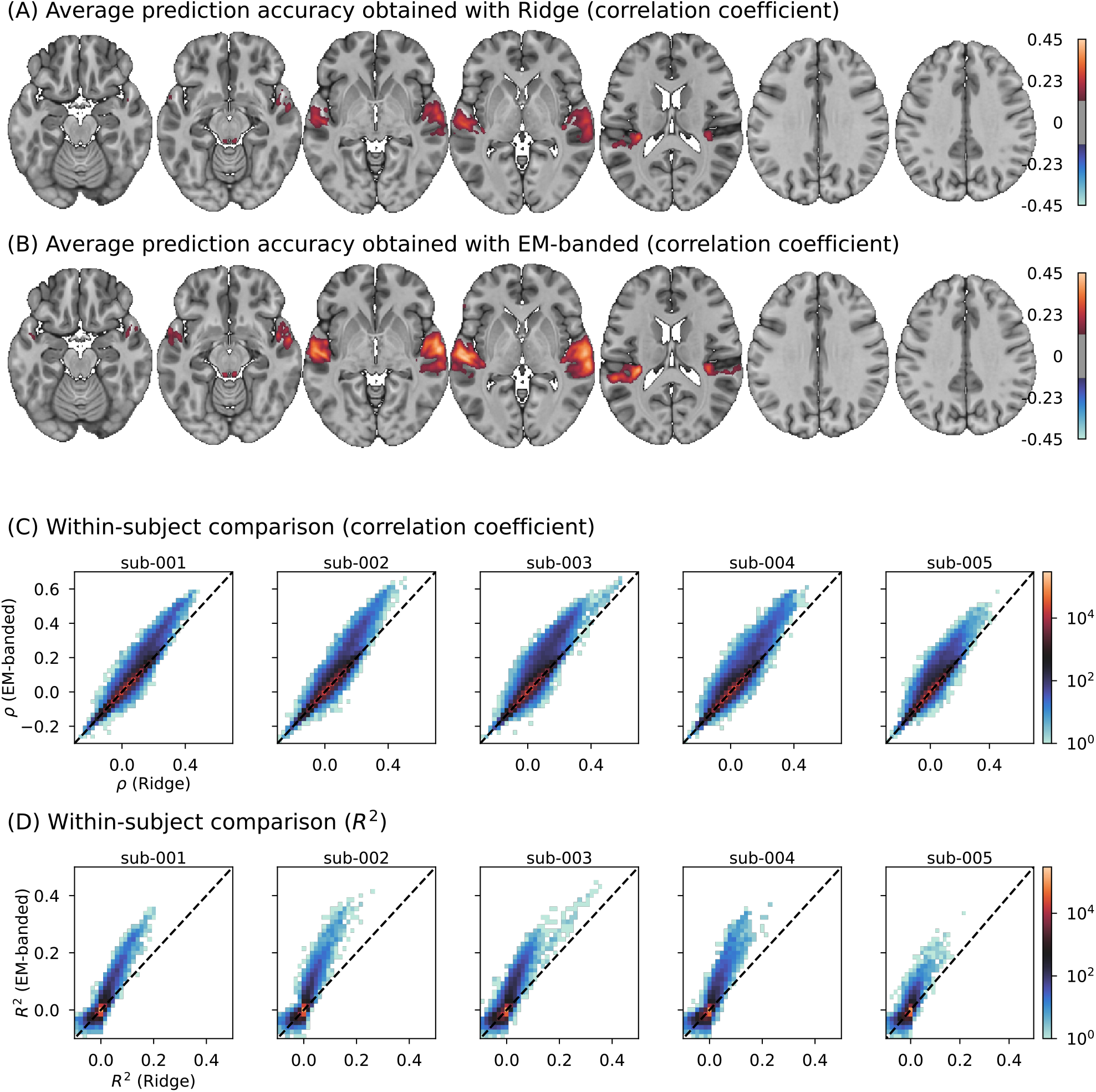
A BOLD fMRI encoding analysis where stimulus-irrelevant predictors are included in the analysis. (A): group-mean prediction accuracies for models trained with the EM-banded framework. (B): group-mean prediction accuracies for cross-validated Ridge regression models. Maps in (A) and (B) have been arbitrarily thresholded for visualization purposes. (C) 2D histogram visualization of prediction accuracies (Pearson correlation coefficient, *ρ*) for the EM-banded and Ridge estimators for each subject. The dashed line indicates similar performance across models. (D) Similar visualizations as in (C) but using *R*^2^ as performance metric.

The regularization term *α* of the Ridge estimator varies substantially across voxels. This can occur, for instance, if voxels with higher signal-to-noise ratio require less overall regularization. Changes in *α* impacts the overall scale of all weights. This can lead to misleading spatial weight maps indicating effects of predictors that are stimulus-irrelevant. This is illustrated in figure 8 showing the group-mean standard deviation of predictions obtained with stimulus-irrelevant predictors. The predictions were obtained using only the task-irrelevant predictors to predict voxel responses (discarding the spectrogram and genre features) with the Ridge estimator. As can be seen, the standard deviation of these predictions is high specifically in auditory cortical regions. This may reflect that the Ridge estimator declares less overall shrinkage in voxels in regions with higher SNR. In effect, the map can misleadingly be interpreted to suggest regional auditory-evoked activation to a random feature *because of* the regularization. Encouraging differential shrinkage of each feature space with the EM-banded estimator avoids this problem, showing no clear spatial patterns in these maps.

**Figure 8.**
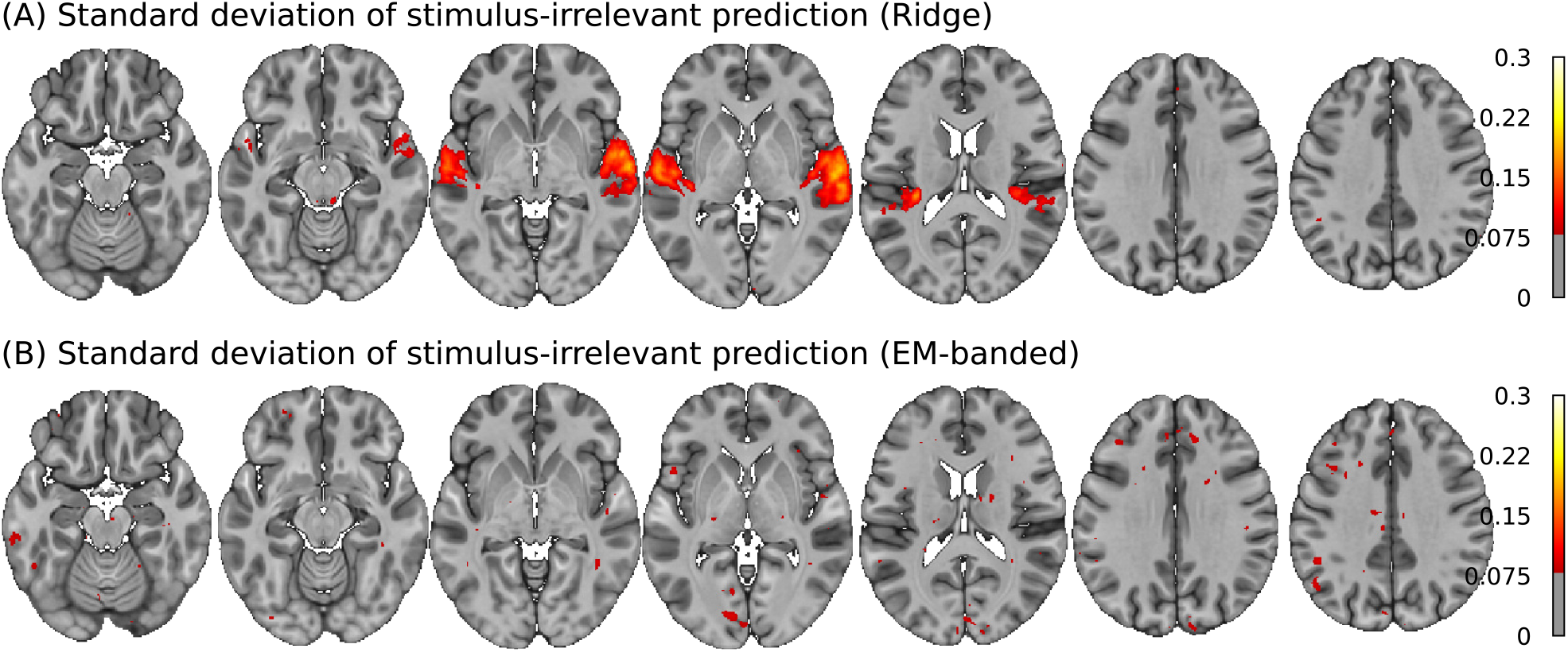
Group-mean standard deviations of predictions with a stimulus-unrelated noise feature estimated with Ridge regression (above) and the EM-banded framework (below).

## 5 EEG-based decoding analysis

In this example, we consider a decoding analysis of single-trial EEG responses to continuous natural speech. The example follows a popular approach of predicting the continuous speech amplitude envelope using linear combinations of time-lagged EEG scalp electrode responses at lower frequencies (Ding and Simon, 2012; O’Sullivan et al., 2014; Wong et al., 2018; Crosse et al., 2016; Broderick et al., 2018). In this approach, the goal is to find a multi-dimensional filter *w* that linearly maps from (features in) the multi-channel EEG response to the envelope of the speech stimulus *y*. The decoding analysis can be formulated as *y*(*t*) = ∑_*c*_ ∑_*τ*_ *x*(*t* −*τ, c*)*w*(*c, τ*) + *ϵ* where *x*(*t* −*τ, c*) denotes preprocessed electrode response in channel *c* at time point *t* − *τ* and where *τ* indicates the time delay between stimulus and response. The model can be specified by concatenating multiple time-lagged versions of each EEG electrode response in a design matrix *X* in Eq. 1. The regression problem now amounts to estimating coefficients of a multi-dimensional FIR filter (de Cheveigné et al., 2018). In this context, it is natural to consider each time-lagged electrode response as a feature group *F*_*j*_ in Eq. 2 and allow for differential shrinkage of weights associated with each group. For instance, it can be relevant to allow for excessive shrinkage of a noisy or seemingly irrelevant electrode for all its time-lag copies.

While EEG speech decoding studies often focus on low-frequency activity (*<* 15 Hz), it is known from intracranial recordings that the power of higher frequency neural activity in the gamma range also track features of speech stimuli (Mesgarani and Chang, 2012; Pasley et al., 2012; Crone et al., 2001). These frequencies are typically highly attenuated in scalp EEG. Nonetheless, including high gamma power features has been shown to improve predictive accuracy of EEG decoding models for some subjects (Synigal et al., 2020). However, including more EEG features, some of which may have very low SNR, is also be associated with a risk of adding predictors to the decoding model that introduce noise to the predictions. It may thus be desirable to allow for differential shrinkage of different EEG features. In this example, we therefore explore predictive accuracy of the EM-banded model when both lower and higher frequency EEG features are included in such a decoding analysis. This example uses publicly available EEG data (https://doi.org/10.5061/dryad.070jc) described in Broderick et al. (2018). In brief, 19 subjects listened to an audiobook while continuous EEG data from 128 scalp electrodes were recorded. Data were acquired in 20 trials each approximately 180 s in length. We fit decoding models again using the EM-banded framework compared to standard Ridge regression. For both models, we used time-lagged low-frequency (LF) EEG features (1-8 Hz) as well as time-lagged higher-frequency (HF) EEG power features (25-35 Hz) (Giraud and Poeppel, 2012) to predict a speech envelope feature. Further details related to EEG preprocessing, audio feature extraction, and model fitting are described in section 2 in the Supplementary Material.

Note that cross-validation was only used for hyperparameter tuning in the Ridge model, but that both models always were tested on held-out data.

The results from the analysis are shown in figure 9. The prediction accuracies obtained with the EM-banded estimator are modestly but consistently higher than the Ridge estimates (*p <* 0.01, Wilcoxon signed rank test). The average correlation between the predicted speech envelope and the target envelope is 0.1368 ± 0.0383 for the EM-banded model and 0.1267 ± 0.0.0377 for the Ridge model. The EM-banded approach excessively shrinks groups of weights, whereas the Ridge estimator yields a denser distribution of weights. Figure 9 (right) shows an example of weights estimated by the two models for one subject. This illustrates how the overall scale of weights associated with the HF features and LF features are comparable for the Ridge estimator whereas the EM-banded estimator differentially shrinks both weights associated with HF features and groups of LF features associated with single electrodes. Given that high-frequency EEG features are weak and in some subjects not even measurable by scalp electrodes, including HF features can negatively model impact predictions, in particular with a Ridge estimator. Differential shrinkage of groups of weights may in such cases allow for better exploitation of potentially informative HF features.

**Figure 9.**
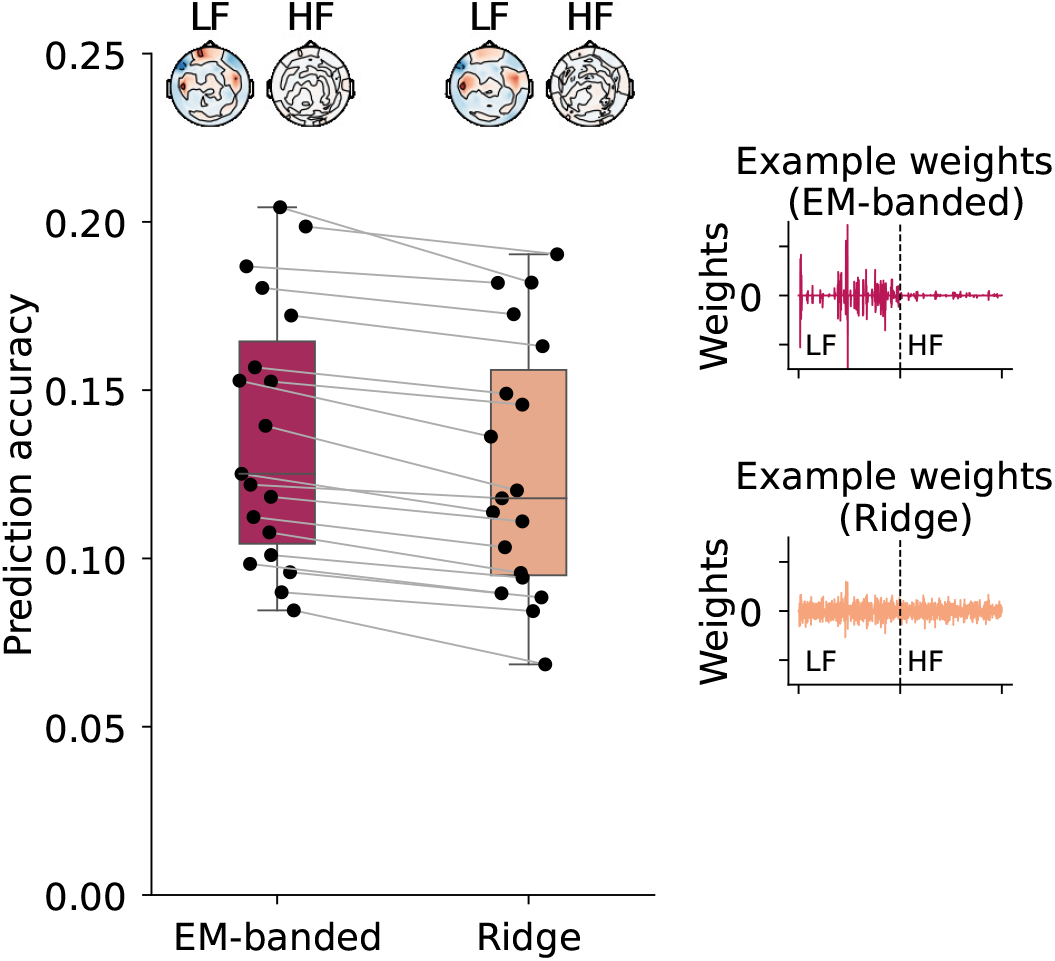
Results from the EEG decoding analysis. Left: boxplot of prediction accuracies obtained with each model. Each point reflects data from a given subject. Prediction accuracy was defined as the average correlation coefficient between the target envelope and decoded envelope in held out trials. Topographies depict weights of each model at a single time lag for illustrative purposes. The weights are here first normalized by their standard deviation and averaged across participants. Topographies are shown both for weights associated with low-frequency (LF) features and higher-frequency (HF) features. Right: Examples of regression weights for one participant.

## 6 Discussion

We explored an empirical Bayes framework for estimating ”banded”-type encoding- and decoding models. The framework can allow differential shrinkage of groups of weights associated with different groups of predictors (e.g., feature ”bands”). The model can further be used to encourage smoothness on such groups of weights. In the following, we discuss modeling considerations and model limitations.

### 6.1 Encoding- and decoding models with correlated predictors

Regression models with highly correlated predictors frequently occur in encoding- and decoding analyses. Features of continuous naturalistic stimuli (e.g., video or speech) used as predictors in encoding models are often highly correlated. Continuous scalp EEG channels used as predictors in decoding models are typically also highly correlated, and pairs of channels can commonly exceed *ρ* = 0.8 with high-density EEG recordings.

The described estimator may encourage solutions where subsets of correlated predictors are excessively shrunken. However, the true posterior of weights in analyses with many correlated predictors may show complex multimodal distributions, and point estimates of model parameters (or, approximate conditional distributions) should be interpreted with caution (see e.g., Simulation 5). Spurious excessive shrinkage of a given feature set in an encoding analysis with correlated feature sets could, for example, be misinterpreted as indicating that the feature is not encoded in a response where it is contributing weakly. Such issues can be particularly difficult to identify when there are many outcome variables (e.g., different voxels with different noise profiles), and little training data to support the identification of model parameters.

Shrinkage procedures can complicate interpretation of weights in encoding analyses with correlated predictors, as illustrated in our simulations. In decoding models, model prediction accuracy on held-out data is often a primary measure. If the main goal is to optimize model performance for some application (e.g., a decoding model optimized for BCI performance), then the choice of shrinkage procedure may be less critical for interpretation as long as it does not compromise performance or lead to exclusion of important features. However, another goal may be to correlate prediction accuracies with behavior or clinical outcome measures. Here, the choice of shrinkage procedure can affect group-level inferences. For instance, multi-channel EEG data may show distinct noise characteristics in different individuals (e.g., noisy electrodes or attenuation of higher-frequency activity due to head anatomy). A given shrinkage procedure might differentially impact decoding accuracies across individuals, potentially complicating the interpretation of group-level correlations with behavior.

With highly correlated predictors, it can sometimes be relevant to transform (groups of) the predictors using data dimensionality reduction techniques (de Cheveigné and Simon, 2008; de Cheveigné and Parra, 2014; Hyvärinen and Oja, 2000; Zou et al., 2006; de Cheveigné et al., 2019; de Cheveigné, 2021) and then fit models using the transformed data. This can potentially reduce computational burden without compromising predictive accuracy. Dimensionality reduction can be combined with banded regularization procedures to allow differential shrinkage of regression weights associated with each component. The described framework can also be used in the context of feature selection (Fan and Lv, 2008; Bair et al., 2006; Neal and Zhang, 2006; Guyon and Elisseeff, 2006), for instance, to inform about which stimulus-response lags to consider. Whether the assumptions entailed by such procedures are reasonable again depends on the specific research goals.

Several of our simulations assumed that there is a set of ”true” regression weights as defined by data-generating equations. For example, in Simulation 4 we assumed that only one of two correlated feature groups drives a response. Such data-generating processes are deliberately over-simplisitic in order to illustrate properties of the model, but are unlikely to reflect realistic scenarios. In naturalistic scenarios, correlated data are likely to be generated by a chain of latent causes. For example, in encoding analyses with different sets of highly correlated features (e.g., different high- and low-level features derived from natural speech), it is conceivable that the different stimulus features as well as the target regression variables each are generated by one of more underlying variables unknown to the modeller. Therefore, regression weights do not allow straightforward causal interpretation in such scenarios, and banded regression procedures do not solve such issues.

### 6.2 Computational burden

Encoding- and decoding analyses often involve high-dimensional regression problems. In encoding models, the stimulus or task feature selection is often motivated by some pre-defined hypotheses. Yet, even for well-defined hypotheses, feature sets often translate to a high number of predictors. This, in turn, implies a high computational burden. The proposed EM-banded model involves matrix inversions (often with complexity 𝒪 (*D*^3^) unless specific structure of the matrix can be exploited) which may limit the relevance if the number of predictors is very high. This problem is further amplified in situations with many response variables. Algorithms described in la Tour et al. (2022); Nunez-Elizalde et al. (2019) are well-optimized for fMRI encoding analyses with many predictors and a large number of response variables (voxels). For this situation, section 3 in the Supplementary Material presents an alternative formulation of the EM-banded model where it is assumed that the *λ*_*j*_ and *ν* parameters are shared across multiple outcome variables. This formulation allows for a more efficient implementation in scenarios with many outcome variables. Runtime test examples can be found at https://github.com/safugl/embanded.

### 6.3 Risks of under-shrinkage and over-shrinkage

The described model applies independent inverse-gamma priors on *λ*_*j*_ terms associated with different groups of predictors. The hyperpriors could also be omitted and replaced by a constant which will lead to typical type-II maximum likelihood estimates. Several efficient algorithms have specifically addressed such scenarios (Tipping and Faul, 2003; Wipf and Nagarajan, 2007; Cai et al., 2021). The choice of hyperpriors, or lack hereof, can have a large impact on the results and misspecified hyperpriors can be associated with over-shrinkage or under-shrinkage, as exemplified in our simulations. In both cases, this may compromise both predictive accuracy and interpretability of model weights.

In the encoding and decoding analysis examples, we focused on inverse-gamma hyperpriors with hyperparameters fixed to *τ* = *η* = *κ* = *ϕ* = 10^−4^, which corresponds to broad hyperpriors. Choosing broad hyperpriors is convenient in several practical analysis situations, allowing for excessive shrinkage of (groups of) predictors while also putting prior mass on larger values for *λ*_*j*_ and *ν*. Moreover, fixing hyperparameters allows for determination of *λ*_*j*_ and *ν* terms from training data alone without having to tune hyperparameters using cross-validation. This can be particularly convenient in banded regression problems with a many feature groups. Section 7 in the Supplementary Material shows out-of-sample predictive accuracies for EM-banded models with various different hyperpriors fit to data from three of our simulations. Here, the predictive accuracies tend to plateau as *γ* approaches zero, suggesting that broad hyperpriors are reasonable choices in these simulations.

In many practical applications, a broad hyperprior can be chosen to provide an initial ’default’ reference model that can be used as a starting point for subsequent model comparisons (Gelman, 2006). In the context of encoding- and decoding analyses (where focus mostly is on out-of-sample predictive accuracy) such reference models can be highly useful for identifying cases where certain regularization properties are inappropriate. This procedure could be taken, for example, when exploring whether grouping structures or smoothness constraints are relevant. The EM-banded model offers one estimator among several others that can be considered in such a process.

### 6.4 Empirical Bayes and related work

We explored a parametric empirical Bayes framework (Morris, 1983; Efron and Morris, 1975, 1973; Robbins, 1964; MacKay, 1992, 1996) in the context of encoding- and decoding analyses. We used this framework to tune regularization hyperparameters from the data and subsequently focus on moments in a conditional distribution, an idea that is widely adopted in hierarchical analyses of neuroimaging data as outlined by Friston et al. (2002b). Parametric empirical Bayes has been considered in the context of MRI decoding analyses (Sabuncu and Van Leemput, 2011; Wen et al., 2019), in M/EEG source localization problems (Owen et al., 2012, 2008; Friston et al., 2008; Wipf et al., 2010; Cai et al., 2021, 2018), in multi-subject hierarchical fMRI analyses (Friston et al., 2002a), in MRI segmentation (Iglesias et al., 2015; Puonti et al., 2016, 2020) and in receptive field models of cell recordings (Park and Pillow, 2011). Likewise, dividing variables into groups and encouraging group-wise sparsity via the group-lasso (Yuan and Lin, 2006) has also been used in M/EEG source localization problems (Haufe et al., 2008; Ou et al., 2009; Lim et al., 2017) and in causal discovery (Haufe et al., 2010; Tank et al., 2017; Bolstad et al., 2011). The group-lasso is convenient in these latter situations because it promotes truly (group-wise) sparse solutions unlike the EM-banded model and banded Ridge regression (la Tour et al., 2022; Nunez-Elizalde et al., 2019), where weight pruning (such as setting excessively shrunken regression weights to zero) would otherwise be necessary.

It is important to once again stress that the considered a modeling framework may provide poor approximations to the true posterior distribution over the model weights. The hope is instead that the model yield reasonable out-of-sample predictions (Tipping, 2001), which we indeed found to be the case in several analyses. In encoding or decoding analyses, the emphasis is typically on out-of-sample predictive accuracy and point estimates of model parameters. Our motivation behind highlighting model weights in several of our simulations was to better illustrate model properties and highlight regression problems where group-shrinkage can be relevant.

### 6.5 Over-fitting and cross-validation

Tuning hyperparameters using cross-validated estimates of expected prediction accuracy may reduce the risk of over-fitting. One benefit of tuning hyperparameters using cross-validation compared to empirical Bayes (without cross-validation) is that it optimizes an estimate of expected prediction accuracy which can lead to good generalization, especially in cases with many observations, high SNR, and independent training and validation splits. Moreover, such procedures can be highly useful for identifying model misspecifications. However, with limited and noisy data, the variance of the expected prediction accuracy estimate can be high. Tuning many free hyperparameters to optimize such an estimate can be associated with a considerable risk of over-fitting in these circumstances.

Cross-validation can similarly be used to tune hyperparameters of the EM-banded model. This may prove beneficial in some cases (compared to fixing model parameters a priori) and could potentially reduce the risk of over-fitting, but the same concerns apply here. In practice, rather than tuning all free hyperparameters of the EM-banded jointly, one may choose to introduce the constraint that only *γ* is tuned, with *γ* = *τ* = *η* = *κ* = *ϕ*. This will practically make it more straightforward to fit EM-banded models across a range of *γ* parameters and tune *γ* to minimize expected prediction error on held-out data. Section 7 in the Supplementary Material shows results from such procedures.

### 6.6 Other perspectives

We used the described modeling framework to fit encoding- and decoding models trained on data from single subjects (i.e., single-subject analyses). The described framework could also be incorporated in “searchlight” decoding analyses (Kriegeskorte et al., 2006) where different subsets of features from multi-dimensional neural data recordings are used as predictors in single-subject or group-level decoding models. The framework could for example be relevant in studies that attempt to use one or more features extracted from multiple imaging modalities to predict clinical scores (Woo et al., 2017). One appealing aspect of group shrinkage procedures is that they can allow for ranking importance of feature groups. This may aid interpretation of encoding analyses (la Tour et al., 2022; Nunez-Elizalde et al., 2019). In recent years, encoding models have been used increasingly to model neural responses with features extracted from task-optimized neural networks (Kell et al., 2018; Tuckute et al., 2023; Yamins and DiCarlo, 2016; Yamins et al., 2014). This typically involves very high-dimensional feature sets and hence encoding models where the number of predictors is much larger than the number of observations, i.e., *D* ≫ *M*. Surprisingly, over-parameterized models may show good generalization performance even with vanishing explicit regularization in minimum-norm least squares estimators (Belkin et al., 2019; Kobak et al., 2020; Hastie et al., 2022). This behavior is clearly different from cases where explicit regularization is critical for performance. Yet, the ability to establish importance of sets of features can similarly be useful in these settings, and group shrinkage procedures may be one way of achieving this.

## 7 Conclusion

In this paper, we explored a framework for estimating banded-type regression models in encoding- and decoding analyses. It offers a tool that can have relevance for specific modelling problems in computational neuroscience, complimenting other related estimators. We used simulations and data examples to illustrate properties and limitations of the described model in comparison with Ridge estimators.

## Funding

This work was supported by the Center for Auditory Neuroscience grant from the William Demant foundation and by a synergy grant from Novo Nordisk foundation to the UHEAL project Uncovering Hidden Hearing Loss (grant number NNF17OC0027872). HRS was supported by a grand solutions grant “Precision Brain-Circuit Therapy - Precision-BCT)” from Innovation Fund Denmark (grant number 9068-00025B) and a collaborative project grant “ADAptive and Precise Targeting of cortex-basal ganglia circuits in Parkinson’s Disease - ADAPT-PD” from Lundbeckfonden (grant number R336-2020-1035). OP was supported by a grant from Lundbeckfonden (grant number R360–2021–395). KHM was supported by the Pioneer Centre for AI, DNRF grant number P1.

### Acknowledgements

The authors would like to thank Edmund C. Lalor (University of Rochester) for helpful discussions.

## Data and Code Availability

Code implementations in both Matlab (The MathWorks, Inc., Natick, Massachusetts, United States) and Python are available on https://github.com/safugl/embanded. The Python code utilizes libraries such as *Numpy* (Harris et al., 2020), *Scipy* (Virtanen et al., 2020) and *Pytorch* (Paszke et al., 2019). The public repository contains examples and runtime tests. EEG data and BOLD fMRI data used in this study have been presented previously (Broderick et al., 2018; Nakai et al., 2022) and is available on https://doi.org/10.5061/dryad.070jc and https://openneuro.org/datasets/ds003720, respectively.

## 1 fMRI encoding analysis

### 1.1 fMRI preprocessing

We used a publicly available BOLD fMRI dataset that has been described in (Nakai et al., 2022, 2021) which contains data acquired from five participants (procedures were approved by local ethics and safety committees (Nakai et al., 2021, 2022)). We considered a minimal BOLD fMRI preprocessing pipeline which involved motion correction (Friston et al., 1996), spatial smoothing (with a 3 mm FWHM Gaussian kernel), and brain extraction. These processing steps utilized SPM 12.6685 (Penny et al., 2011) and FSL 6.0.1 (Jenkinson et al., 2012; Smith et al., 2004). Part of the motivation for incorporating spatial smoothing was to improve signal-to-noise ratio, but also that susceptibility-induced distortions rendered it difficult to adequately align volumes across functional runs. Six motion regressors (translations in x, y, and z directions and rotations about x, y, and z) were extracted for each subject and functional run. These were used to create a set of nuisance regressors that further included a cosine basis set from a one-dimensional discrete cosine transform. Time courses from each voxel were deflated using this set of regressors. We defined the average response to a given sound stimulus clip as the average of these residualized voxel time courses from 6 s to 16.5 s after stimulus onset. This was a pragmatic choice and it means that only a subset of each stimulus clip contributes to the estimated ”average” response (while minimizing contamination from the previous stimulus clip). For the last stimulus clip, we averaged from 6 s after stimulus onset to the last acquired volume. The initial 15 s of ’dummy’ images in each functional run were omitted from subsequent analyses in both test and training runs. All analyses were conducted in native subject space. For visualization purposes, we mapped the resulting maps of predictive accuracy to a Montreal Neurological Institute (MNI) template. This was achieved using tools available with Advanced Normalization Tools (ANTs) (Avants et al., 2009). Visualization of results relied on scientific computing software in Python, and utilized libraries such as *Nilearn* (Abraham et al., 2014), *Seaborn* (Waskom, 2021), *Numpy* (Harris et al., 2020), and *Matplotlib* (Hunter, 2007).

### 1.2 Feature groups

We defined three features groups, two of which were related to the audio stimuli. The first, *F*_1_, was a genre-label dummy-coded feature that indicated which of 10 genres each stimulus clip had been assigned to by Tzanetakis and Cook (2002). The second, *F*_2_, was a time-averaged ”spectrogram” representation. To generate this representation, each stimulus clip was passed through a 32-channel gammatone filterbank (Slaney et al., 1993) with filter centre frequencies spaced between 50 Hz and 11050 Hz (Glasberg and Moore, 1990). Envelopes were extracted from the output of each filter via the absolute value of the Hilbert transform and subsequently power-law compressed with a compression factor of *c* = 0.3. We defined the spectrogram representation of each stimulus clip as the temporal average of these features from 2 s to 11 s after stimulus onset. Finally, we considered a third feature group, *F*_3_, which contained 400 stimulus-irrelevant noise predictors. These predictors were simulated for each stimulus block as a random vector from a multivariate Gaussian distribution with zero mean and covariance *C* = exp(−0.3|*k* − *i*|^2^), *k* = 1, …, 400, *i* = 1…400.

### 1.3 Regression

Separate regression models were fit to data from each voxel. Predictors and the voxel response time courses were standardized. For each voxel, we fit models to data from the 12 training runs and evaluated predictive accuracy on data from held-out test runs. For the EM-banded estimator, we defined distinct *λ*_*j*_ parameters to each of the three feature groups and defined that *τ* = *η* = *κ* = *ϕ* = 10^−4^. For the Ridge estimator, we considered a 5-fold cross-validation procedure to inform the selection of *α* from data from the 12 training runs. A range of possible *α* values over a grid of 50 logarithmically spaced values between 10^−5^ and 10^10^ were considered. The chosen *α* parameter was the parameter that minimized the average mean square error between model predictions and target responses in held-out folds. Prediction accuracies on the test set were quantified in two ways. First, we computed the Pearson correlation coefficient between predicted responses in test runs and the target responses. Second, for each voxel time course, we computed *R*^2^ defined as follows:

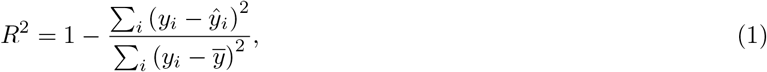

where *ŷ*_*i*_ is the predicted response, where *y*_*i*_ is the target response, and where 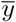 is the mean response.

## 2 EEG decoding analysis

### 2.1 Preprocessing

We used a publicly available EEG dataset https://doi.org/10.5061/dryad.070jc which includes EEG data from 19 participants that each undertook 20 experimental trials in which they listened to speech stimuli (Broderick et al., 2019) (procedures were approved by local ethics committees). The dataset includes a ’broadband’ envelope representation of the speech stimulus in each experimental trial. The first-order temporal difference of this envelope representation was computed and the output was then half-wave rectified (Synigal et al., 2020; Hertrich et al., 2012; Daube et al., 2019), centered and band-pass filtered between 1 Hz and 8 Hz using a second-order Butterworth filter. Zero-phase filtering was achieved by filtering in the forward and reverse directions. The dataset includes EEG data from each experimental trial sampled at 128 Hz. These data were re-referenced to the common average, detrended (second-degree polynomial trends), and low pass filtered at 40 Hz using a fourth-order zero-phase Butterworth filter. We defined two EEG feature groups from these preprocessed data: low-frequency (LF) features and higher-frequency (HF) features. For the LF features, we filtered the data between 1 Hz and 8 Hz using the same filters as those applied to the audio envelope representations. For the HF features, we filtered the data between 25 Hz and 35 Hz using second-order zero-phase Butterworth filters and then computed the absolute value of the Hilbert transform from each electrode time course. These time courses were subsequently filtered between 1 Hz and 8 Hz using again the same filters as those applied to the audio envelope representations. Finally, the EEG and audio features were downsampled to 40 Hz to reduce computation time. We augmented EEG features from each channel with a set of time lags by concatenating the time-lagged features side-by-side so as to absorb temporal mismatches between stimulus features and EEG features in the subsequent regression (De Cheveigné et al., 2021). The time lags ranged from -250 ms to 0 ms. We focused on data from 10 s post-trial onset to approximately 160 s post-trial onset for both EEG and audio features.

### 2.2 Regression

We focused on decoding models that utilized time-lagged LF and HF EEG features to predict the speech envelope feature. We used a leave-one-trial-out outer cross-validation approach, fitting models to data from all trials except one and evaluating prediction accuracy on the held-out trial. This procedure was repeated for all trials. For the EM-banded model, we grouped predictors into *j* = 1, …, 256 sets, each consisting of multiple time-lagged versions of either an LF or HF feature from a single electrode. During cross-validation, we computed the means and standard deviations of the training data. Subsequently, we transformed both the training data and the held-out data by subtracting these mean values and dividing by these standard deviations. This was done separately for each predictor and for the target variable. The scale of features extracted from single-trial EEG can vary dramatically across subjects and electrodes, and it can thus be convenient to standardize the data when focusing on default prior distributions on model weights and default initialization schemes.

For the EM-banded model, we fixed hyperparameters related to the Inverse-Gamma priors to *γ* = 10^−4^, and further encouraged smoothness on temporal filters for each predictor group by defining that *h*_*j*_ = 0.5 for all *j* = 1, …, 256 groups (with *h*_*j*_ defined as in the main text). For the Ridge estimator, we further considered an inner 5-fold cross-validation procedure. The *α* parameter was tuned to maximize average correlation between predictions and target envelope features in held-out inner folds. We considered *α* values ranging from 10^−5^ to 10^10^ in 50 logarithmically spaced steps.

Prediction accuracies were defined as the Pearson correlation between model predictions and target envelope feature in held-out test trials. The correlation coefficients were averaged across trials after first applying the Fisher-Z transform and the average was inverse-transformed using the inverse of the Fisher-Z transform. Visualization of results in Python utilized libraries such as *MNE-Python* (Gramfort et al., 2013), *matplotlib* (Hunter, 2007) and *seaborn* (Waskom, 2021).

## 3 Multiple output case

Here we formulate an EM-banded model for the regression scenario with multiple outcome variables and a design matrix *X* that is shared for all outcome variables. Let *Y* ∈ R^*M*×*P*^ be a matrix with *p* = 1, …, *P* outcome variables stacked column-wise. In this case, the regression problem can be written in matrix notation as follows:

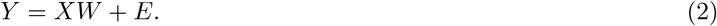

Here, *X* ∈ R^*M*×*D*^ denotes the design matrix, *W* ∈ R^*D*×*P*^ denotes the weights and *E* ∈ R^*M*×*P*^ denotes error terms. We will once again assume that the design matrix *X* can be partitioned into *J* meaningful groups such that *X* = [*F*_1_, …, *F*_*j*_, …, *F*_*J*_], where each group *F*_*j*_ has one or more predictors stacked column-wise. Let *D*_*j*_ denote the number of predictors in the *j*-th group such that the total number of predictors equals *D* = ∑_*j*_ *D*_*j*_. Further, let 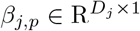 be a column vector that contains *D*_*j*_ weights associated with predictor group *j* for outcome variable *p*. We partition *W* and Λ as follows:

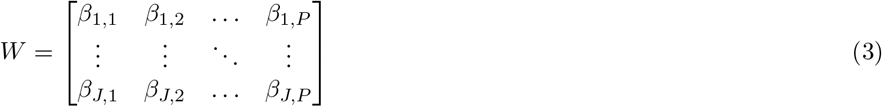

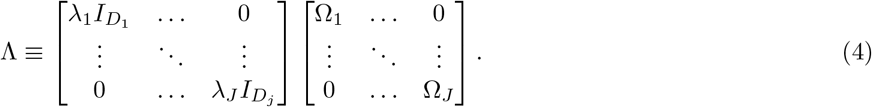

Each Ω_*j*_ block has size *D*_*j*_ × *D*_*j*_ and follows the definition in the main text. We will henceforth represent vectorized versions of *Y* and *W* as *y* and *w* respectively, defined as follows:

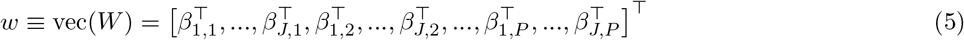

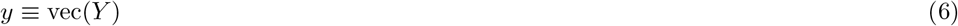

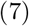

where vec(*Y*) denotes the vec-operator applied on *Y*. Additionally, let *H* ≡ *I*_*P*_ ⊗ *X* where ⊗ denotes Kronecker product. To simplify notation, we will again let *λ* denote a set of 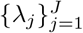 parameters and not highlight dependence on terms that remain fixed. We will assume that each outcome variable has been scaled in a way that justifies the following model:

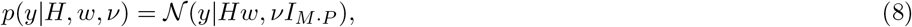

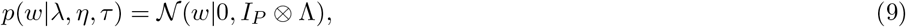

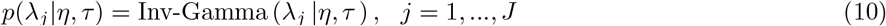

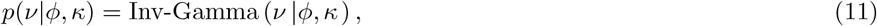

Here, *I*_*M* ·*P*_ denotes an identity matrix of size (*M* · *P*) ×(*M* · *P*). Letting Σ ≡ (Λ^−1^ + *ν*)^−1^*X*^⊤^*X* ^−1^ and *B* ≡ *ν*^−1^Σ*X*^⊤^*Y* we see that:

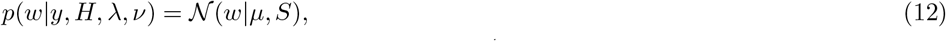

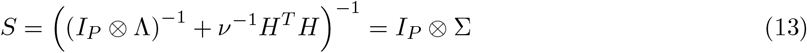

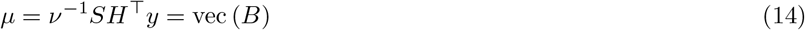

We let 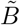 and 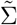 denote *B* respectively Σ estimated for a given set of *λ* and *ν*. We subsequently partition the matrix 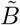 as follows:

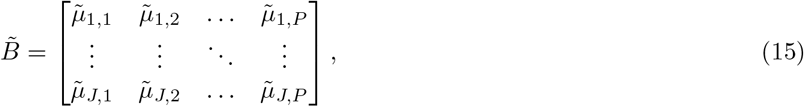

where 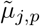 denotes a column vector with elements associated with predictor group *j* for outcome *p*. We let 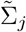 denote a block of size *D*_*j*_ × *D*_*j*_ along the diagonal in 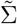 associated with group *j*. We follow the same procedure as in the main text and find the following closed-form update rules:

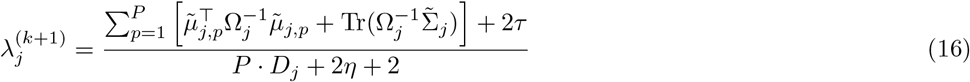

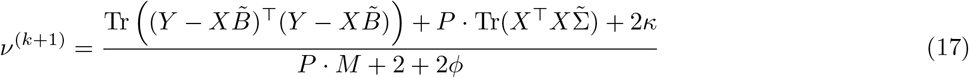

To simplify this expression further, we define that 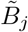 is a block in 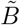 of size *D*_*j*_ × *P* with elements associated with group *j*, such that 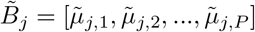. We can now write:

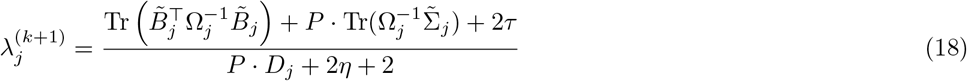

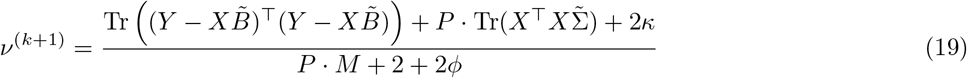

It should be clear that the above model is similar to the model described in the main text when *P* = 1. Note that it can be relevant to compute *X*^⊤^*X* and *X*^⊤^*Y* and avoid computing these terms multiple times. It can also be relevant to use the cyclic property of the trace in some scenarios (e.g., when *P > D*). Additionally, it can be relevant to make use of the Woodbury matrix identity (Murphy, 2012) and define Σ = Λ − Λ*X*^⊤^(*νI*_*M*_ + *X*Λ*X*^⊤^)^−1^*X*Λ when *D* ≫ *M*.

### 3.1 Log-objective

We recall that our goal is to maximize the marginal posterior density *p*(*λ, ν*|*y, H*) which is equivalent to maximization of *p*(*λ*)*p*(*ν*)*p*(*y*|*λ, ν, H*). Here, the logarithm of *p*(*y*|*λ, ν, H*) takes the form:

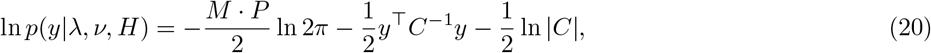

with *C* = *νI*_*M*·*P*_ + *H*(*I*_*P*_ ⊗ Λ)*H*^⊤^. The matrix *C* has size (*M* ·*P*) × (*M* ·*P*) which could be problematic if one were to directly evaluate Eq. 20 in a regression problem with many outcome variables. We follow Tipping (2001) and simplify this expression using the Woodbury matrix identity and the matrix determinant identity. In this case, ln |*C*| can be simplified and written as follows:

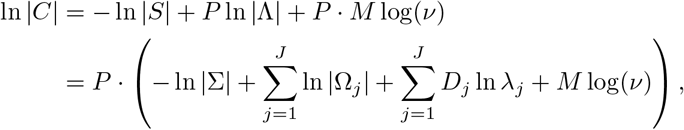

Notice that Σ has size *D* × *D*. One can similarly simplify *y*^⊤^*C*^−1^*y* and express it in terms of *B*:

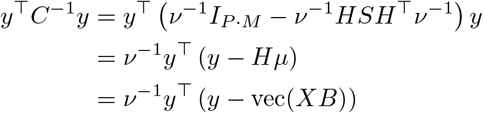

We can now write the expression in Eq. 20 as follows:

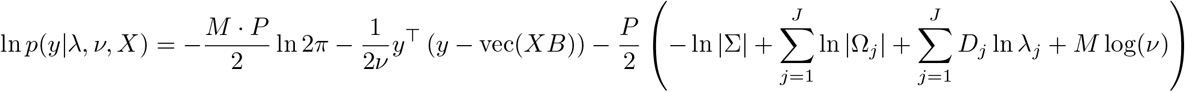

This is usually fast to compute even when there is a high number of outcome variables and hence dimensions in *C*. Our implementation of the algorithm incorporates this expression when computing the logarithm of the objective function (hereafter referred to as log-objective) at each iteration. Ignoring terms that remain fixed during optimization with a fixed set of hyperprior parameters, we express the log-objective Δ as follows:

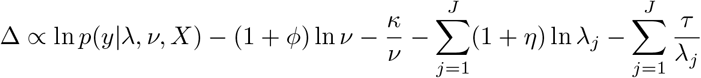

The implementation makes it possible to define a convergence criterion based on increases in the log-objective, such that the algorithm will terminate if increases in the log-objective are below some user-defined tolerance. We leave it up to future studies to explore alternative (faster) approaches to maximizing the log-objective.

## 4 Simulating responses to acoustic and phonetic features

### 4.1 Feature groups

Data from the *The DARPA TIMIT Acoustic-Phonetic Continuous Speech Corpus* (TIMIT) (Garofolo et al., 1993) were used for this simulation example. We utilized sentences spoken by 190 speakers for this example, with each speaker delivering ten sentences. Audio waveforms were root mean square normalized. Auditory spectrogram features were extracted from each audio waveform. To generate these features, we passed each waveform through a 32-channel gammatone filterbank (Slaney et al., 1993) with filter centre frequencies spaced between 50 Hz and 8000 Hz. Envelopes were extracted from the output of each filter via the absolute value of the Hilbert transform and subsequently power-law compressed with a compression factor of *c* = 0.3. The envelopes were lowpass filtered at 64 Hz (second-order zero-phase Butterworth filter) and downsampled to 128 Hz. For the phonetic features, we defined boxcar regressors based on time-aligned phonetic labels available from each sentence (Garofolo et al., 1993; Zue and Seneff, 1996). Specifically, we constructed six boxcar regressors flagging time periods where the annotation indicated the occurrence of the following categories: stops, affricatives, fricatives, nasals, semivowels and glides, and vowels. We define these boxcar regressors as phonetic features in this simulation. It should be stressed that our goal is not to make statements about the coding of phonetic features, but rather to illustrate properties of regularized estimators when considering such stimulus feature sets. Spectrogram features and phonetic features were extracted from each sentence. Next, we created multiple time-lagged versions of each predictor in these feature sets for the time lags: 0, 1, …, 38. The first 39 samples of features extracted from each sentence were subsequently discarded. This procedure was repeated for all sentences and the features were stacked row-wise and subsequently standardized. We let *F*_1_ denote time-lagged spectrogram features and *F*_2_ denote time-lagged phonetic features. The procedure resulted in a total of 661633 samples, corresponding to approximately 86 minutes of data. The feature group *F*_1_ has 1248 columns and the feature group *F*_2_ has 234 columns

### 4.2 Alternative data-generating process

Simulation 4 in the main text assumes that the response variable only contains a mixed version of *F*_1_, but not of *F*_2_. For completeness, Figure 1 depicts results of a similar simulation where the response is simulated as *y* = *F*_2_*w*_2_ + *ϵ*, such that only the phonetic features are assumed to affect the response. Ridge models and EM-banded models are again fit to the data using both feature sets. The simulated ”target” weights *w*_2_ are shown in Figure 1A.

**Figure 1.**
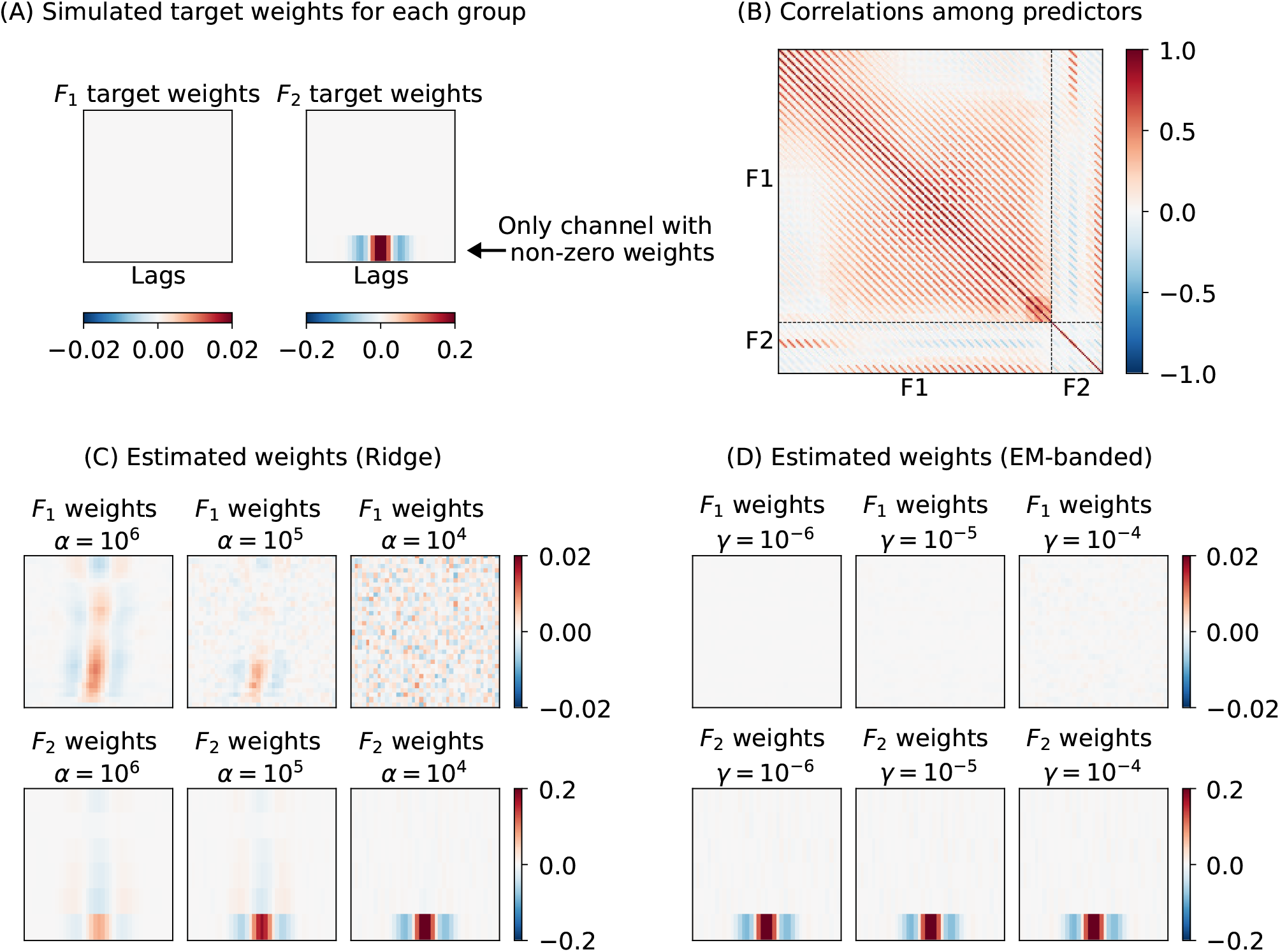
(A) Target weights associated with each of the two feature sets. Here, only phonetic features are simulated to affect the response. (B) Matrix of Pearson correlation coefficients between predictors. Dashed lines are used to visualize predictor groups, *F*_1_ and *F*_2_. (C) Weights estimated with Ridge estimator for each of the two groups. (D) Weights estimated with the EM-banded estimator for each of the two groups. Ridge estimators with *α* set to 10^6^, 10^5^ and 10^4^ are shown. EM-banded estimators were considered with *γ* set to 10^−6^, 10^−5^ and 10^−4^

## 5 Simulation with groups of correlated predictors

Here, we consider a simulation in which we simulate two groups of predictors, *F*_1_ and *F*_2_. Each of these two feature groups has 128 dimensions and 2048 rows. We assume *y* = *F*_1_*w*_1_ + *ϵ* and that only *F*_1_ affects the outcome variable, *y*. The target weights *w*_1_ are shown in Figure 2. As previously, we assume that *ϵ* is drawn from a Gaussian distribution. Each row in [*F*_1_, *F*_2_] is drawn from a multivariate Gaussian with zero mean and a covariance *C* defined as follows:

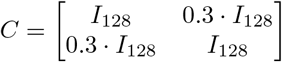

This simulates correlations among predictors associated with the different groups. The SNR is 0 dB. We again compare Ridge regression model fits with EM-banded model fits using a set of fixed hyperparameters. The estimated weights for the Ridge and EM-banded model are shown in Figure 2. The EM-banded estimator excessively shrinks the weights associated with *F*_2_ (as desired) when *γ* attains a low value. In all cases, the EM-banded model accurately recovers the weights associated with *F*_1_. The Ridge estimator, on the other hand, tends to pull correlated weights towards each other when *α* attains a high value (in this case when *α* = 1000 and *α* = 10000). This has the undesired consequence that the weights associated with *F*_2_ tend to resemble the weights associated with *F*_1_ more. This could potentially lead to misinterpretation of the weights associated with *F*_2_.

**Figure 2.**
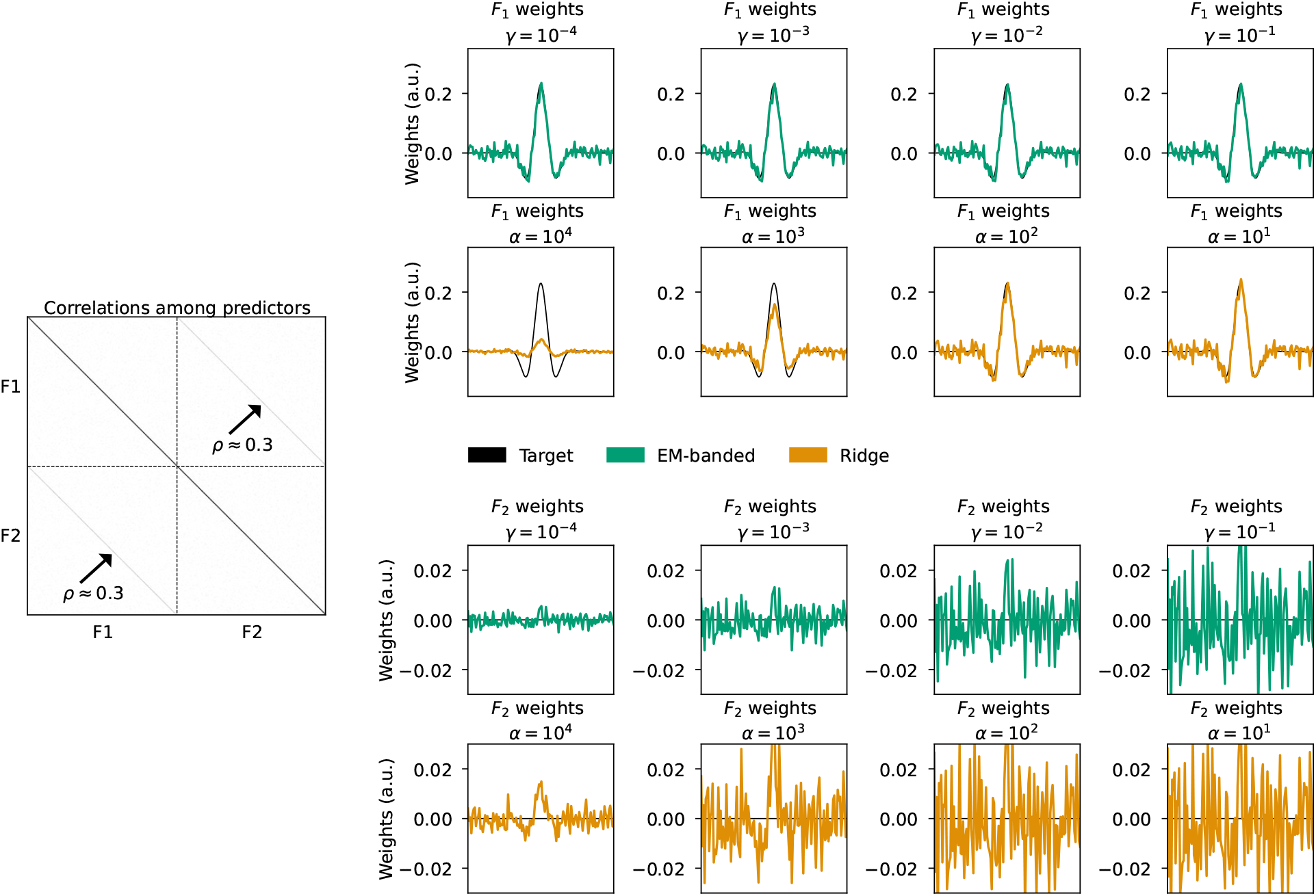
Simulation 6 illustrating behavior of the Ridge estimator and the EM-banded estimators when there are correlations among predictors that have been divided into two groups, *F*_1_ and *F*_2_. Left middle panel: Matrix of Pearson correlation coefficients between predictors. Dashed lines are used to visualize predictor groups, *F*_1_ and *F*_2_. Right: Weights estimated with Ridge estimator and with the EM-banded estimator. The top two rows indicate weights associated with *F*_1_. The bottom two rows indicate weights associated with *F*_2_. The target weights are shown in black.

## 6 Banded regression and hyperparameter tuning

Banded Ridge regression avoids making explicit assumptions about the prior distribution of hyperparameters and instead uses cross-validation to tune hyperparameters. Using cross-validation to tune regularization strength and optimize predictive accuracy on held-out data may be a strategy to facilitate feature selection and interpretability of model parameters. However, tuning multiple free hyperparameters using cross-validation can also lead to spurious excessive shrinkage of weights associated with a given feature band, e.g., if the variance of the expected prediction accuracy estimate is high, complicating the interpretation of model parameters.

To illustrate this point, the following simulation shows an unfavorable SNR situation similar to Simulation 5 in the main text, but here focuses on a banded Ridge regression estimator. As in Simulation 5, we simulate two predictors, *F*_1_ and *F*_2_, and a response variable *y* = *F*_1_*w*_1_ + *F*_2_*w*_2_ + *ϵ*. The predictors, as well as the noise term *ϵ*, are drawn from Gaussian distributions, and the number of samples is 512. The target weights are fixed to non-zero values and the SNR is approximately −20 dB. We now focus on the following estimator:

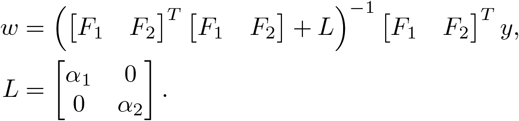

This estimator is fit to the training data for different values of *α*_1_ and *α*_2_. When *α*_1_ = *α*_2_, this corresponds to a standard Ridge estimator.

Next, we simulate five independent validation sets, each with 512 samples. The simulation process for predictors, noise, and target variables mirrors that of the training set. For example, for the first validation set, we simulate two predictors 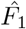 and 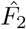 and response variable 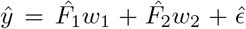, where 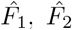, and 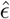 are drawn from Gaussian distributions, and where *w*_1_ and *w*_2_ are the same target weights as for the training set. For each validation set, we evaluate how well the model predicts the target variable *ŷ* for different values of *α*_1_ and *α*_2_.

If one were to select hyperparameters, *α*_1_ and *α*_2_, to minimize MSE (or maximize correlation), then this may lead to excessive shrinkage applied to one of the two weights, to neither of the weights, or to both, depending on the validation set. Moreover, the level of regularization declared to each weight may strongly depend on performance metric used for parameter tuning (e.g., MSE). Using similar tuning procedures for the Ridge estimator may also result in different amounts of shrinkage depending on the specific validation set, but in this case, (nearly) the same amount of shrinkage would be applied to both weights (hence having little impact on Pearson correlation coefficients).

These simulations are highly stylized and mainly serve to assist intuition about how interpretability of model parameters in banded regression with multiple ’bands’ similarly can be complicated, also when regularization hyper-parameters are tuned using cross-validation. This once again highlights the relevance of exploring various models to better understand whether model properties are inappropriate for a given problem.

**Figure 3.**
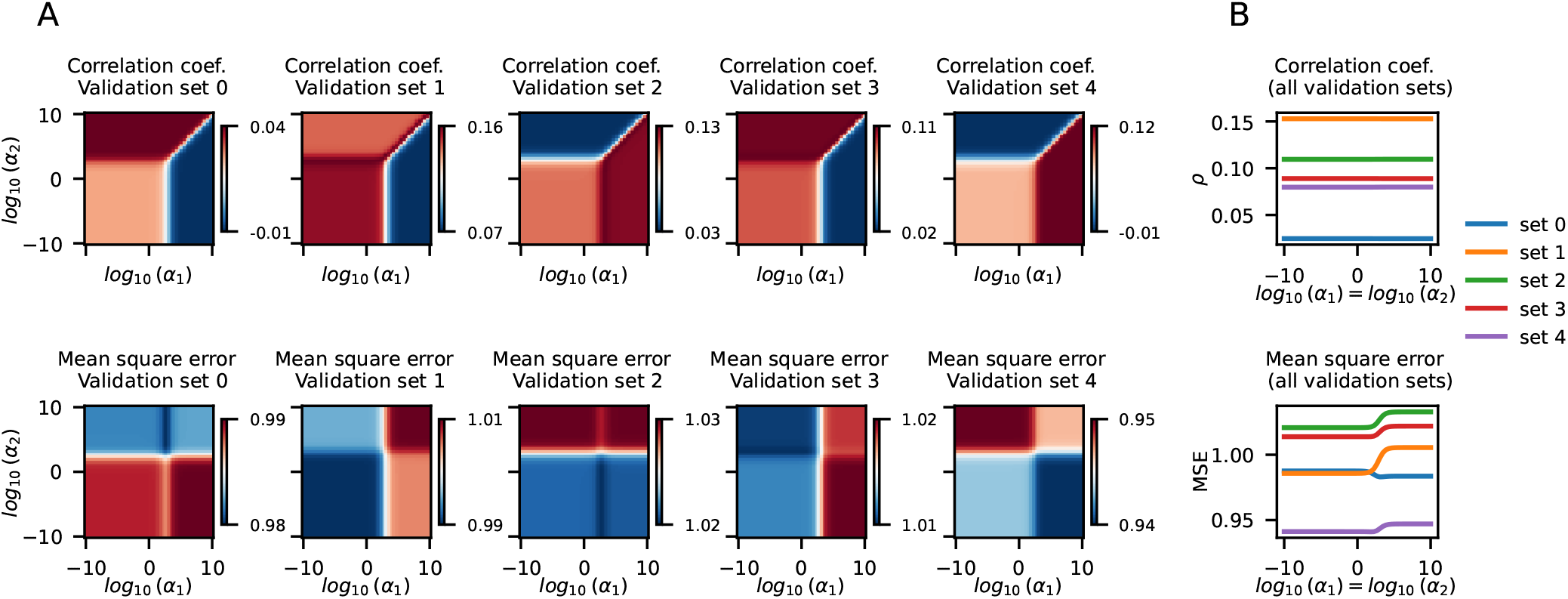
Results from a simulation with a single training set and multiple validation sets. (A) Results from analyses across a range of *α*_1_ and *α*_2_ values. (B) Results from analyses with *α*_1_ = *α*_2_. Each line in panel (B) represents data from a single validation set. Top row: Pearson correlation coefficient between model predictions and target variable in each validation set. Botton row: Mean square error between model predictions and target variable in each validation set.

## 7 Out-of-sample prediction accuracy

Simulations 1-3 in the main text illustrate weights estimated by the EM-banded model in different simulation scenarios. Here, we explore how well the models in these simulations predict novel data with different hyperparameter settings. For each of these three simulations, we simulate validation data using the same data-generating processes as for the training data. The same target weights are considered for both the training data and the validation data. For instance, in Simulation 1, we simulate three groups of predictors (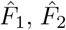, and 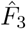, each with 64 predictors) and simulate a response variable *ŷ* as 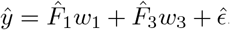, where *w*_1_ and *w*_3_ represent the same target weights as described in the main text, and where 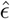 indicates Gaussian noise. We now fit EM-banded models to the training data and attempt to predict *ŷ* given the model and given 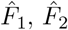, and 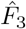. We evaluate predictive accuracy in two ways: using mean square error between model predictions and *ŷ*, and using the Pearson correlation coefficient between model predictions and *ŷ*. This procedure is repeated for all three simulations and for EM-banded models fit with different values of *τ, η, κ*, and *ϕ*. Additionally, the procedure is repeated for Ridge regression models. For completeness, we also illustrate prediction accuracy across a range of *a* and *b* parameters where we define:

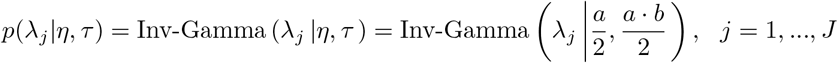

Results from all these analyses are shown in Figures 4, 5, and 6 for Simulations 1, 2, and 3 respectively. Notice that these figures also show results from analyses where we introduce the constraint that *τ, η, κ*, and *ϕ* all should equal the same value, *γ*. We observe that prediction accuracy tends to plateau as *γ* attains low values in all these simulations (see Fig. 4C, Fig. 5C, and Fig. 6C).

**Figure 4.**
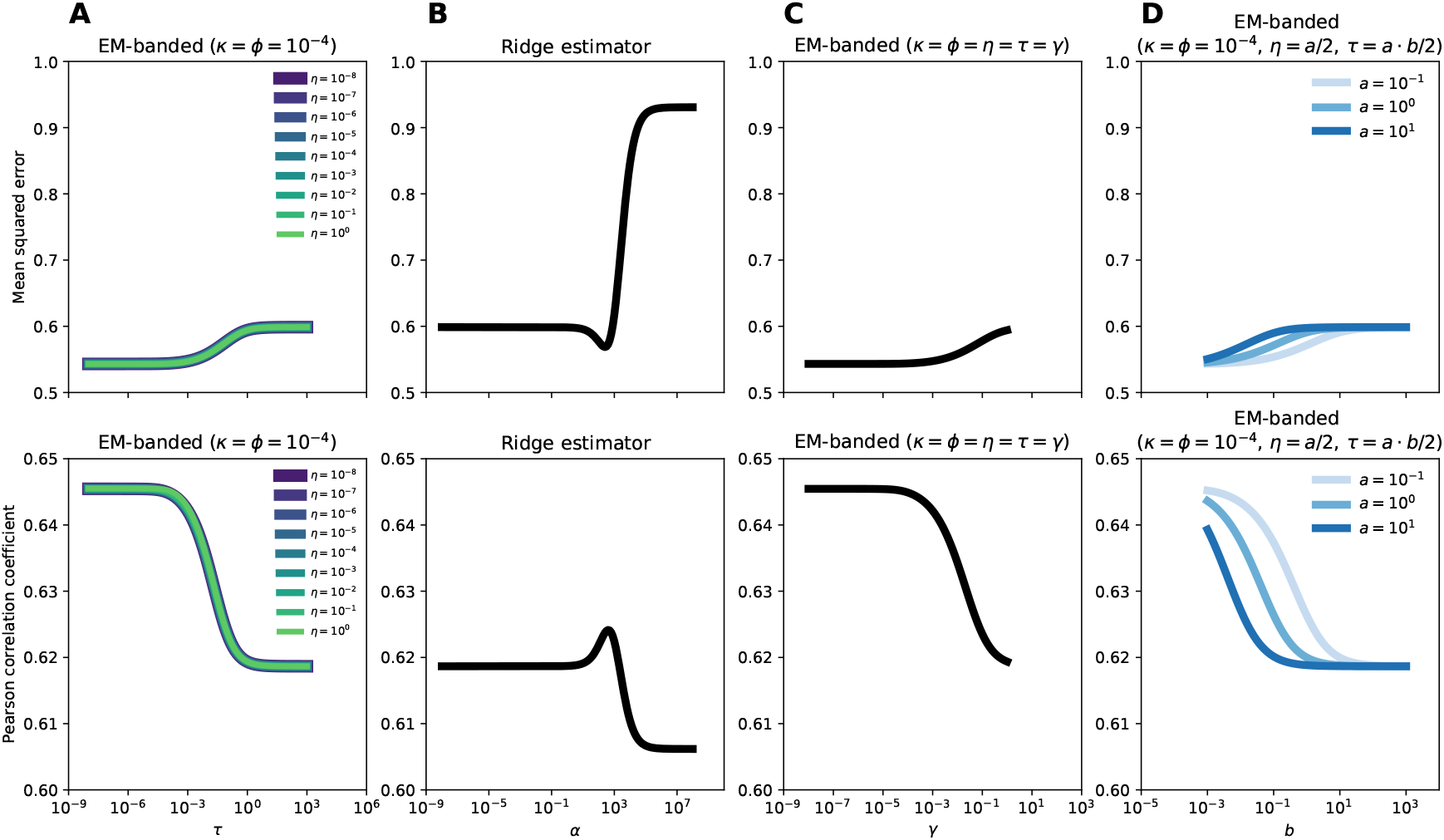
Simulation 1. Top row: Mean square error between model predictions and target variable. Bottom row: Pearson correlation coefficient between model predictions and target variable. (A): results obtained with EM-banded models fit with different values for *τ* and *η*, but with *κ* and *ϕ* kept fixed to a low value of 10^−4^. (B): results obtained with the Ridge model across a range of *α* parameters. (C): results obtained with EM-banded models fit with different values for *γ* where *τ* = *η* = *κ* = *ϕ* = *γ*. (D): results obtained with EM-banded models fit with different values for *a* and *b* where *η* = *a/*2 and *τ* = *a* · *b/*2, and where *κ* and *ϕ* are kept fixed to 10^−4^.

**Figure 5.**
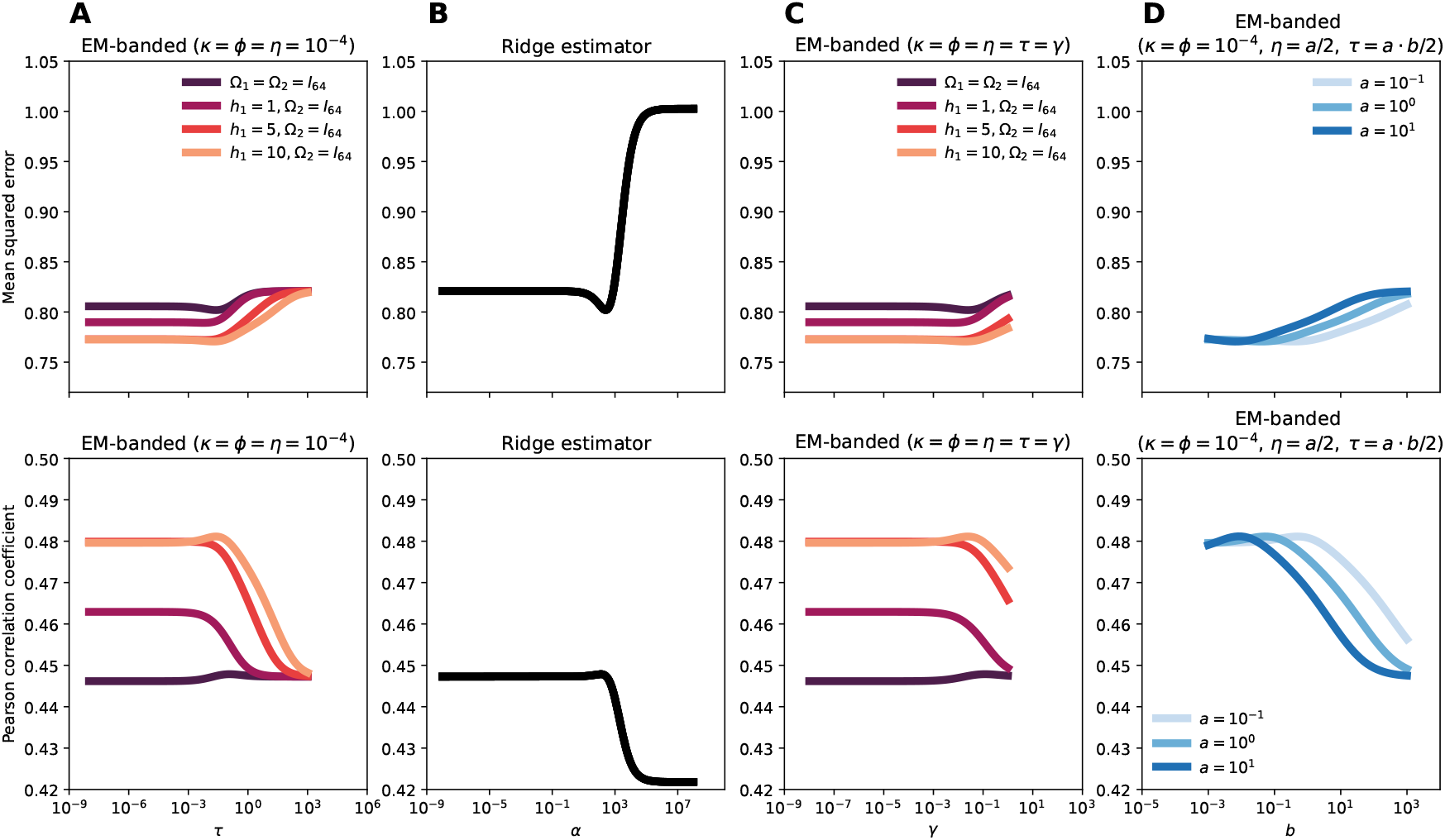
Simulation 2. Top row: Mean square error between model predictions and target variable. Bottom row: Pearson correlation coefficient between model predictions and target variable. (A): results obtained with EM-banded models fit with different values for *h*_1_ and *τ*, but with *κ, η* and *ϕ* kept fixed to a low value of 10^−4^. (B): results obtained with the Ridge model across a range of *α* parameters. (C): results obtained with EM-banded models fit with different values for *γ* and *h*_1_ where *τ* = *η* = *κ* = *ϕ* = *γ*. (D): results obtained with EM-banded models fit with different values for *a* and *b* where *η* = *a/*2 and *τ* = *a* · *b/*2. Here, we kept *κ* and *ϕ* fixed to 10^−4^ and further kept *h*_1_ fixed to 10.

**Figure 6.**
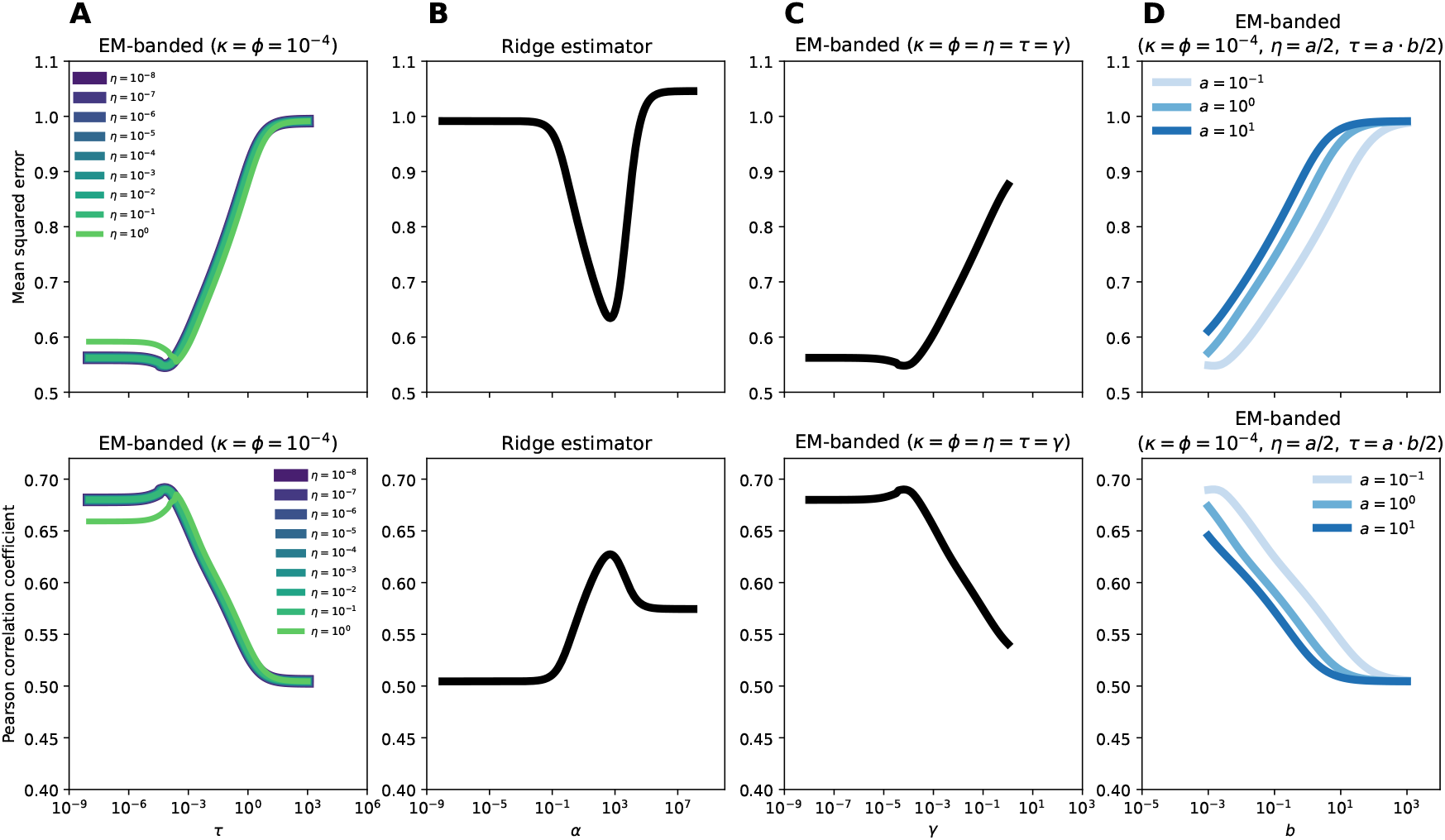
Simulation 3. Top row: Mean square error between model predictions and target variable. Bottom row: Pearson correlation coefficient between model predictions and target variable. (A): results obtained with EM-banded models fit with different values for *τ* and *η*, but with *κ* and *ϕ* kept fixed to a low value of 10^−4^. (B): results obtained with the Ridge model across a range of *α* parameters. (C): results obtained with EM-banded models fit with different values for *γ* where *τ* = *η* = *κ* = *ϕ* = *γ*. (D): results obtained with EM-banded models fit with different values for *a* and *b* where *η* = *a/*2 and *τ* = *a* · *b/*2, and where *κ* and *ϕ* are kept fixed to 10^−4^.

